# GEM-DeCan: Improved tumor immune microenvironment profiling through novel gene expression and DNA methylation signatures predicts immunotherapy response

**DOI:** 10.1101/2021.04.09.439207

**Authors:** Ting Xie, Jacobo Solórzano, Miguel Madrid-Mencía, Abdelmounim Essabbar, Julien Pernet, Mei-Shiue Kuo, Alexis Hucteau, Alexis Coullomb, Nina Verstraete, Olivier Delfour, Francisco Cruzalegui, Vera Pancaldi

## Abstract

Quantifying the proportion of the different cell types present in tumor biopsies remains a priority in cancer research. So far, a number of deconvolution methods have emerged for estimating cell composition using reference signatures, either based on gene expression or on DNA methylation from purified cells. These two deconvolution approaches could be complementary to each other, leading to even more performant signatures, in cases where both data types are available. However, the potential relationship between signatures based on gene expression and those based on DNA methylation remains underexplored.

Here we present five new deconvolution signature matrices, based RNAseq data or on DNA methylation, which can estimate the proportion of immune cells and cancer cells in a tumour sample. We test these signature matrices on available datasets for in-silico and in-vitro mixtures, peripheral blood, cancer samples from TCGA, and a single-cell melanoma dataset. Cell proportions estimates based on deconvolution performed using our signature matrices, implemented within the EpiDISH framework, show comparable or better correlation with FACS measurements of immune cell-type abundance and with various estimates of cancer sample purity and composition than existing methods.

Using publicly available data of 3D chromatin structure in haematopoietic cells, we expanded the list of genes to be included in the RNAseq signature matrices by considering the presence of methylated CpGs in gene promoters or in genomic regions which are in 3D contact with these promoters. Our expanded signature matrices have improved performance compared to our initial RNAseq signature matrix. Finally, we show the value of our signatures in predicting patient response to immune checkpoint inhibitors in three melanoma cancer cohorts, based on bulk tumour sample gene expression.

We also provide GEM-DeCan: a snakemake pipeline, able to run an analysis from raw sequencing data to deconvolution based on various gene expression signature matrices, both for bulk RNASeq and DNA methylation data.

## Background

The tumor microenvironment (TME) is defined as the collection of cells and extracellular matrix that surround cancer cells inside a tumor. It affects tumor development through interactions between the different cells. This impacts the probability for cancer cells to escape immune-control, grow and metastasize, and plays an important role in therapy response and resistance (DeBerardinis, 2020).

Much of the recent progress in cancer treatment derives from the exploitation and reactivation of immune cells, such as lymphocytes, that infiltrate the TME and fight cancer cells. Despite the great potential of immuno-oncology, there is a considerable difference in efficacy of these therapies across tumor types and patients. It is thus of paramount importance to develop tools to identify the different types of immune cells present in biopsy samples, as this could aid personalized therapy approaches.

More specifically, recent findings about the importance of myeloid cells in hampering the response to immunotherapies make the development of macrophage signature matrices an important goal for immuno-oncology (Engblom et al., 2016). CD8+ T cells are normally responsible for killing cancer cells through their cytotoxic activity. Traditional immunotherapies rely on the concept that these killer CD8 cells are already infiltrated in the tumor region, where their interaction with the cancer cells can disable their cytotoxic activity (O’Donnell et al., 2019). Unfortunately, a high percentage of patients (Sharma et al., 2017) do not seem to respond durably to these treatments and recent findings implicate some types of myeloid cells in the TME of these patients in these response failures.

In particular, tumor-associated macrophages (TAMs) can be found in the microenvironment of solid tumors in high numbers. Depending on their phenotypes, TAMs can promote tumor progression, by suppressing antitumor immunity, or directly protecting cancer cells (Anderson et al., 2017) and they are important treatment targets in cancer, especially in tumors which do not present high lymphocyte infiltration (Mantovani et al., 2017; Pathria et al., 2019).

TAMs can be differentiated from circulating monocytes upon entering the tumor area or pre-existent tissue resident cells and they acquire different phenotypes that can be indicatively distributed along a spectrum of polarization states going from M1 polarization, promoting inflammation, to M2, with a pro-tumor character (Lawrence & Natoli, 2011). However, these cells are extremely plastic and our knowledge on their behaviour comes from in-vitro experiments that are not fully representative of conditions inside tumors (Lawrence & Natoli, 2011; Murray, 2017).

Since macrophages are not present in blood and have very plastic phenotypes, developing fingerprint signatures to identify them from bulk tumour samples has proven challenging. (Lawrence & Natoli, 2011; Murray, 2017).

A common approach to identify and quantify the presence of specific cell types in bulk samples after cell separation from tissues or cultures based on their surface markers is Fluorescence Activated Cell Sorting (FACS). This technique can become cumbersome and multiple algorithms have recently become available to computationally estimate cell type proportions or tumour purity from bulk RNAseq data (Finotello, Rieder, et al., 2019) in complex cellular mixtures (Aran et al., 2015), a procedure generally referred to as deconvolution. Deconvolution methods can be classified as “reference-based” (Houseman et al., 2012) or “reference-free” (Houseman et al., 2014; Zou et al., 2014), depending on whether a specific signature matrix is used to identify the cell types, or clustering is used to directly infer the different cell types present (Visvader, 2011) and can exploit gene expression or methylation data.

DNA methylation (DNAm) profiles are cell-type specific and an excellent alternative to transcriptomes to perform cell-type deconvolution (Titus et al., 2017) since the methylome can be thought of as a record of the cell’s past history and is less affected by transient perturbations of the cell’s environment (Cavalli & Heard, 2019).

In this paper, we exploited a large collection of haematopoietic epigenomes of reference produced by the BLUEPRINT project (Stunnenberg et al., 2016), and other public data sets (**Additional file 1**) to establish a series of novel signature matrices for gene expression and/or methylation based deconvolution approaches and estimation of sample purity.

Specifically, we further hypothesized that genes that are not present in the gene expression-based signature matrix but associated with CpGs that are in the methylation signature matrix, should also be included in the gene-expression deconvolution signature matrix. We use chromatin structure in haematopoietic cells (Javierre et al., 2016) to identify genes whose expression might be impacted by methylation locally and through 3D contacts, and were able to show that our expression-based signature matrix is improved when including these genes.

We further explore the combination of DNA methylation-based and gene expression based deconvolution results, showing that the combination can provide more robust estimates of cellular proportions in the case in which both data-types are available on the same samples.

Whereas the performance of our approach is in most cases comparable to others on benchmark datasets, we note that our signatures for TAMs perform particularly well as biomarkers of response to anti-PD1 treatment in multiple published datasets

Finally, we provide the GEM-DeCan pipeline to the community to process gene expression and DNA methylation data and apply several deconvolution methods using pre-existing or user-generated signature matrices.

## Methods

### RNAseq data processing

The TCGA expression data normalized as Fragments Per Kilobase of transcript per Million (FPKM) were taken from public datasets (Wang et al., 2018). The transcripts per millions (TPM) expression datasets for WB were downloaded from GTEx portal (https://gtexportal.org/home/). 9 purified blood-derived immune cells TPM expression datasets on GRCh38 (**Additional file 1: Table SA**) were collected from the Blueprint project portal (Stunnenberg et al., 2016). The expression value FPKM was first converted to TPM using the following formula:

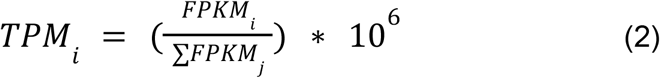

We then merged all the samples together (adjacent non-tumour samples, tumor, WB and immune). The final expression value *TPM_bl_* for each gene *b* in the sample *l* was calculated from TPM with the following formula:

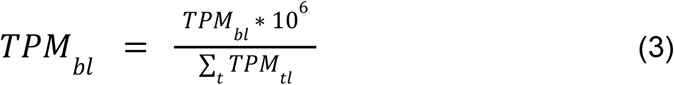

### Development of RNAseq expression signatures for deconvolution

To generate a RNAseq expression signature for cancer cells. We generated first an immune cell type signature using CD4^+^ T cells (N=12), CD8^+^ T cells (N=3), B cells (N=5), Monocytes (N=7), M0 Macrophages (N=4), M1 Macrophages (N=4), M2 Macrophages (N=5), NK cells (N=2) and Neutrophils (N=10) and filtering out T_reg_ cells due to low number of samples (N=1). The files were parsed into R using the *limma* (Ritchie et al., 2015) Bioconductor package and were using the voom function to remove heteroscedasticity by log2(TPM + 1). A series of linear models and empirical Bayes methods were then used to derive the candidate signature features between the given cell types using all pairwise comparisons. The candidate signature features from this analysis were restricted to genes overexpressed that showed a logFC > 2 at a Benjamini-Hochberg corrected p-values < 0.05, with a maximum of 150 genes overexpressed per pairwise comparison **(BPRNA signature, Additional File 3: Table S1)**.

### Generation of the CCLE_TIL10 signature matrix

To develop this signature matrix we started from the TIL10 (170 genes) one proposed in the quanTIseq method (Finotello, Mayer, et al., 2019), which includes 10 immune cell types (B cells, M1 and M2 macrophages, monocytes (Mono), neutrophils (Neu), natural killer (NK) cells, non-regulatory CD4+ T cells, CD8+ T cells, Treg cells, and myeloid dendritic cells (DC)).

FASTQ files of all samples used for generating the TIL10 signature matrix (170 genes) were downloaded, preprocessed and gene expression was quantified as described in (Finotello, Mayer, et al., 2019). An expression matrix, for 10 immune cell types, was constructed, consisting of 19,423 genes and 51 samples. We then constructed a cancer signature matrix (“CCLE”) by considering differential expression between cancer cell line samples from CCLE (eliminating blood cancer cell lines) (Ghandi et al., 2019) and healthy tissues and blood samples from GTEx (GTEx Consortium, 2013). Briefly, we used a bootstrap approach by which 50 samples were randomly taken from each of the three datasets to construct the complete dataset (150 samples in total) for the analysis of differential expression by limma. The mean-variance relationship was modeled with the voom function and the Benjamini-Hochberg method was used for multiple hypothesis testing. lfc > 2.5 and FDR < 0.005 were used to select differentially expressed genes. Only genes which are highly expressed in cancer cells compared to normal tissues and compared to blood samples were selected as “UP” genes. This procedure was repeated 30 times and only the UP genes which are present in each iteration are selected as cancer cell specific genes (**Fig 1a**). The expression profile of cancer cells was computed as the median of the expression values over all samples for all UP genes in the CCLE dataset, resulting in the CCLE signature matrix (138 genes).

**Figure 1:**
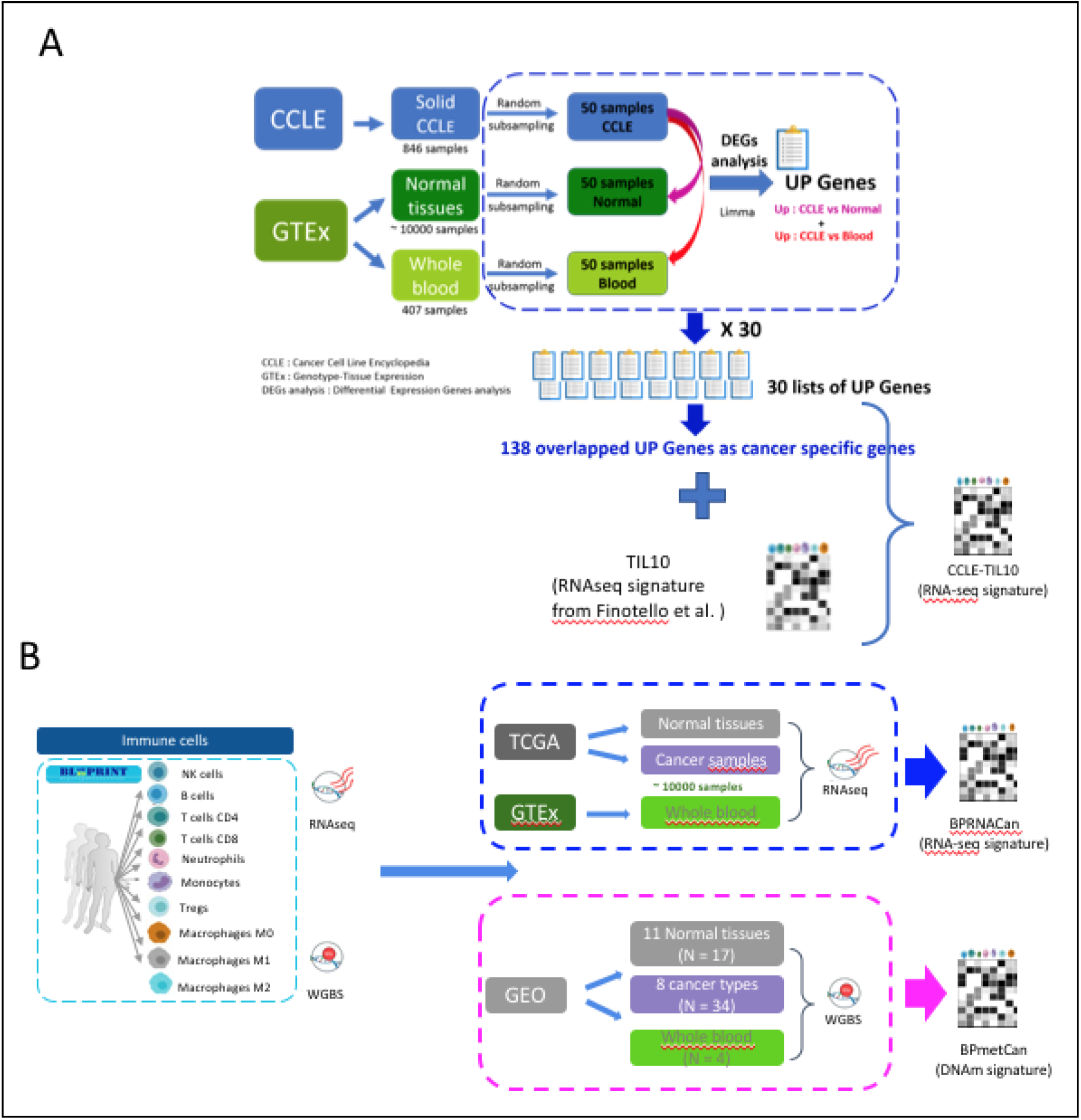
Schematic description of the proposed deconvolution approach. (**a**) To deconvolve the exact proportion of cancer cells as well as immune cells, the TIL10 signature matrix(Finotello, Mayer, et al., 2019) was combined with a list of genes that differs between cancer cell lines, normal tissues and whole blood, generating the CCLE_TIL10 signature matrix. (**b**) Workflow to generate CCLE_TIL10, BPRNACan and BPmetCan signature matrices, through combining TIL10, BPRNA and BPmet immune signature matrices, and cancer signature genes and CpGs (See methods for further details).

To build the combined CCLE_TIL10 signature matrix, we simply considered the union of genes from the TIL10 and CCLE signature matrices (**Fig. 1a**). The expression profiles in the matrix were computed as the median of the expression values over all samples belonging to each cell type (**Additional file 3: Table S2**).

### Generation of the BPRNACan signature matrix

We also built an alternative cancer signature matrix using normal and cancer samples collected from TCGA for different cancer types with normal adjacent tissue and WB from GTEx, using the same procedure as creating the immune cell type signature, selecting logFC > 3 at a FDR < 0.05, with a maximum of 100 genes overexpressed per pairwise comparison. Samples from TCGA were from 18 cancer types: Bladder, Cholangiocarcinoma, Thyroid carcinoma, Liver hepatocellular carcinoma, Colon adenocarcinoma, Kidney Chromophobe, Kidney Renal Clear Cell Carcinoma, Kidney renal papillary cell carcinoma, Lung Squamous Cell Carcinoma, Lung Adenocarcinoma, Stomach adenocarcinoma, Cervical squamous cell carcinoma and endocervical adenocarcinoma, Uterine Corpus Endometrial Carcinoma, Head-Neck Squamous Cell Carcinoma, Breast invasive carcinoma, Rectum adenocarcinoma, Esophageal carcinoma. The final BPRNACan signature matrix was generated by merging BPRNA and this cancer signature, and includes 1403 genes (**BPRNACan Additional file 3: Table S3, Fig. 1b**).

### Collection of WGBS data

To build the Blueprint signature matrix, we collected 52 samples generated from whole-genome bisulfite sequencing (WGBS) of 10 purified blood-derived immune cells on GRCh38 (human genome) (**Additional file 1: Table S2**). Bigwig files including methylation signal and coverage of methylation signal were downloaded from Blueprint epigenome level 3 data (http://www.blueprint-epigenome.eu). In addition, we also downloaded 7 cancer datasets, 4 WB and 11 normal tissues (**Additional file 1: Table S3**) via Gene Expression Omnibus (GEO, https://www.ncbi.nlm.nih.gov/geo/) and Brinkman et al. (Brinkman et al., 2019).

### WGBS data processing

The files were parsed into R data structures, we then discarded bases that have coverage below 10X (Ziller et al., 2015) and also have more than 99.9th percentile of coverage in each sample. Methylation signal (WGBS_*β*_ was calculated with the following formula:

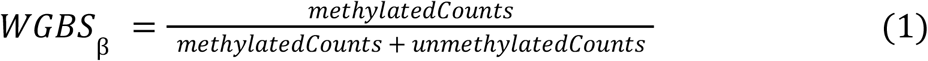

The hg19 coordinates of WGBS from GEO were converted to GRCh38 with the liftOver R package. To map the common methylated cytosines from GEO and Blueprint datasets by genomic position to the Illumina 850K Methylation Array CpG sites, we used the Infinium MethylationEPIC v1.0 B5 Manifest (https://support.illumina.com/downloads). After the coordinate transformation, the missing values on the beta matrix were imputed with impute R package version 1.66.0 (Hastie et al., 2001) to generate the final beta matrix without missing values.

A beta-value matrix was generated from the WGBS dataset, including 409,103 CpGs (48% overlap with EPIC’s 850k CpGs) measured across a total 107 samples (Normal=17, WB=4, Cancer=34, and Immune=52).

### Generation of the DNA methylation signature matrix (BPmet) for blood

We started from WGBS public datasets from the Blueprint project to generate a methylation based signature matrix (BPmet), which we used for immune cell decomposition in blood from healthy donors. Datasets for purified immune cells of 6 types were considered: CD4^+^ T cells (N=8), CD8^+^ T cells(N=8), B cells (N=9), Monocytes (N=4), NK cells (N=2) and Neutrophils (N=7) (Stunnenberg et al., 2016). The signature features only took differentially methylated CpGs (DM-CpGs) using a limma-based wrapper function with a series of linear models and Bayes methods for all pairwise comparisons between candidate cell types that showed an absolute logFC > 2 at an FDR < 0.05, with a maximum of 100 DM-CpGs per pairwise comparison (**BPmet Additional file 3: Table S4).**

### Generation of the enhanced DNA methylation signature matrix for cancer samples (BPmetCan)

To identify cancer cells, normal cells and immune cells in tumour samples, we used firstly a set of Normal (tissues), Whole Blood (WB) and cancer (solid tumours) WGBS samples (Normal: N=17, WB: N=4, Cancer: N=34) via GEO and Brinkman et al. (Brinkman et al., 2019) to generate a signature matrix that can recognize the three groups of cells **(Additional File 1: Table SC)**. To identify the CpGs we needed to include in this signature matrix, we chose CpGs that were highly methylated in cancer cells compared to Normal as well as WB samples with an absolute logFC > 2 at an FDR of 0.05, selecting a maximum of 100 DM-CpGs per pairwise comparison. Specifically In the tumor microenvironment, additional populations of immune cells can be detected compared to blood: macrophages, ranging on a continuum of macrophage polarization that can be exemplified by three macrophage states (M0, M1, M2) (Farha et al., 2020) and T_reg_ cells (Binnewies et al., 2018). For this reason, we extended the BPmet signature matrix with those 4 new immune cell types to generate a new BPmet signature matrix, following the previously detailed signature matrix construction process. We used an absolute logFC > 0.3 at an FDR of 1e-05, with a maximum of 300 DM-CpGs per pairwise comparison.

To build the final BPmetCan signature matrix, which can identify cancer cells and also specific immune types, we merged the above described signature matrix for cancer/normal/blood cells with the BPmet signature matrix extended with macrophages and T_regs_ **(Fig. 1b).** The final significant CpGs are the union of CpGs presented in those two signature matrices, respectively, whereas the profiles of CpGs in each cell type were calculated through the median of methylated values over all samples belonging to that cell type. The BPmetCan signature matrix includes 1896 CpGs (**Additional file 3: Table S5**).

### Generation of gene expression signature matrices expanded according to the methylation signature matrix (BPRNACanProMet) and 3D chromatin contact maps (BPRNACan3DProMet)

Our BPRNACan signature matrix contains 1403 genes which we call *sig genes*, whereas the 1896 CpGs from the BPMetCan signature matrix are denoted as *sig CpGs*. To take into account the potential involvement of genes that are important for each cell type, as evidenced by methylation, but not sufficiently differentially expressed to be *sig genes*, we created a set of expanded gene expression signature matrices.

We considered 3D chromatin contact networks for all immune cells included in Javierre et al. (Javierre et al., 2016), detected by the Promoter Capture Hi-C technique and filtered using CHiCAGO (Cairns et al., 2016). First, we combined BPRNACan *sig genes* with the genes that have a BPMetCan *sig CpG* in their promoter (according to promoter definitions based on promoter capture libraries in (Javierre et al., 2016)), leading to the “BPRNACanProMet” signature matrix (**Additional file 3: Table S6**). Additionally, we generated the “BPRNACan3DProMet” signature matrix by filtering the added genes to take only the ones that have both a *sig CpG* in the promoter and also a 3D contact with a fragment containing a *sig CpG*, (**Additional file 3: Table S7**).

### RNAseq processing and deconvolution pipeline

In order to test the different signature matrices we present in this paper, we provide an RNAseq analysis pipeline, built with snakemake (Köster & Rahmann, 2018) and conda. It allows the user to choose from various tools and options (**Supplementary Fig. S1**). The pipeline can start from raw Illumina sequencing data (.bcl format) or from already processed and normalized TPM data and runs a selection of tools performing deconvolution with the methods used in this paper (quanTIseq and EpiDISH) and all signature matrices mentioned in the paper. It is freely available on the GEM-DeCan repository. To compare our results to those obtained with the CIBERSORTx method (Newman et al., 2019) we ran the algorithm using the default parameters with the Docker image available at https://cibersortx.stanford.edu/download.php upon request.

### Validation datasets

Different datasets were used to test our new signatures, see **Additional file 2: Table S1** for details. Briefly, these included

- in-silico mixtures from (Finotello, Mayer, et al., 2019), consisting of 1700 samples created by in-silico mixing of reads from immune-cells and cancer cell lines in different proportions, simulating different tumor purity (0 to 100%)
- 13 individuals PBMCs with Flow Cytometry estimates: CD4^+^ and CD8^+^ T cells, monocytes, B cells, NK cells (GSE107011, Monaco et al. 2019)
- 19 primary tumor samples from non-metastatic melanoma patients from GSE72056 (Tirosh et al., 2016)
- TCGA Cancer samples with methylation and gene expression profiles as well as cancer and immune cell type experimental estimates. To test the BPmetCan and BPRNACan signature matrices, 16 different TCGA datasets from solid tumors were downloaded in level 3 (beta-value, 450k Illumina array) as well as their corresponding gene expression profiles measured by RNAseq from LinkedOmics (http://linkedomics.org/login.php#dataSource). We added 0.5 to formalize the beta value of methylation data between 0 to 1 due to this methylation being based on zero-centered values.

To estimate the proportion of cancer cells inside tumor samples, a consensus approach integrated four different methods (ABSOLUTE, ESTIMATE, LUMP and IHC) as previously proposed on TCGA samples (Aran et al., 2015) as described in **Additional file 2: Table S2**. Since H&E estimates of cell types only include 2 broad cell lineages (lymphoid, myeloid) and neutrophils, cell types were aggregated as follows:

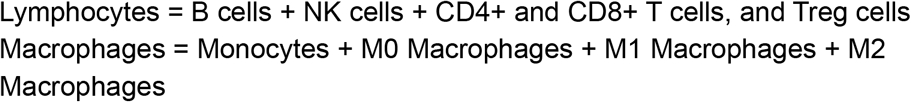

### Whole blood methylation datasets

For validating and assessing the performance of the BPmet signature matrix, we used two independent public datasets (**Additional file 2: Table S1**): 100 WB samples from the Grady Trauma Project (GSE132203) profiled using IlluminaHumanMethylationEPIC and another 6 WB samples using the 450k methylation array from Koestler et al. (GSE77797) (Koestler et al., 2016). Flow-cytometry estimates of the proportion of blood cell types were available for the two WB datasets. The estimated fraction of cells obtained by deconvolution using the EpiDISH (RPC: robust partial correlation) method (Teschendorff et al., 2017) was compared to the flow-cytometry estimated proportions using Pearson Correlation.

### Comparison between proportions estimated by deconvolution and other methods

We used Pearson’s correlation coefficients to compare our estimates of deconvolved proportions to either FACS data or alternatively estimated proportions (**Additional file 2: Table S2**).

### Regression models to predict response to immunotherapy

Three public melanoma datasets with response to anti-PD1 (Gide et al., 2019; Hugo et al., 2016; Riaz et al., 2017) were considered. ElasticNet (Friedman et al., 2010) penalized logistic regression models were run using the results from different deconvolution methods and signature matrices as features.

For each combination of signature matrix and deconvolution method, 4 models were trained, including 3 models trained by leave-one-dataset-out (lodo) and one model trained by 5-fold cross-validation (standard CV). The training includes a hyperparameter search for the l1 ratio and penalty strength. For the lodo training this search is performed by 5-fold CV on training datasets, and models are evaluated on the remaining test dataset. For the standard CV, one fourth of samples is kept as a hold-out test set, and the hyperparameter search is performed by 5-fold CV on the remaining samples, the model is then evaluated on the hold-out test set. Models’ performance were evaluated with the Area Under the Receiver Operating Characteristic Curve (ROC AUC).

## Results

### New immune and cancer cell gene expression signature matrices

To provide an overview of composition of the tumour microenvironment, our main goal was to develop a deconvolution signature that would identify the proportions of immune cells of different types and cancer cells. We started by developing a novel immune cell type deconvolution signature matrix based on RNAseq expression data from primary samples for 6 immune cell types (CD4^+^ T cells, CD8^+^ T cells, Monocytes, B cells, NK cells and Neutrophils) derived from blood (Stunnenberg et al., 2016), which we called BPRNA (see methods, **and Additional file 2: Supplementary text S1, Figures S2a Table S1**). We then proceeded to test the BPRNA signature matrix using two deconvolution methods for which signature matrices can be specified by the user (EpiDISH (Teschendorff et al., 2017) and deconRNASeq (Gong & Szustakowski, 2013)) implemented in our GEM-DeCan pipeline (**Figure S1**) on peripheral blood mononuclear cell (PBMC) samples (**Additional file 2: Table S1)**. Additionally, we benchmarked our signatures against two methods that come with their own signature: MCP-counter (Becht et al., 2016), which is a scoring method based on marker genes, and quanTIseq (Finotello, Mayer, et al., 2019), which is based on constrained least squares regression and can estimate immune cell fractions and fractions of unknown cells with high accuracy (**Additional file 2, Supplementary text S1, Figure S2b**).

Since our main interest is to apply these methods to cancer samples, we set out to extend this initial RNA-based signature matrix for immune cells to detect cancer cells as well (estimating tumour purity). We considered two ways of designing a signature matrix that would estimate tumor purity based on bulk RNAseq expression data, adding Tregs and specific M0, M1 and M2 macrophage signatures to identify the presence of these cell types that are relevant in cancer samples. In the first case, we constructed a signature matrix (CCLE_TIL10, see methods, **Figure 1A** and **Additional file 3: Table S2**) based on RNAseq data for over a thousand cancer cell lines (Ghandi et al., 2019) and a large number of healthy tissue and blood RNAseq samples from the GTEx (GTEx Consortium, 2013).

Aware of the difference between cancer cell lines and cancer cells, we also developed a signature matrix for detecting cancer and specific immune cells starting from expression data in cancer, adjacent normal tissues and immune cells. To this end, we considered samples with RNAseq data for tumor and adjacent non-tumor tissues and whole blood, readily available through TCGA and GTEx, and integrated the resulting signature matrix with the BPRNA immune cell signature matrix. The new integrated signature matrix, which we called BPRNACan, consists of 1403 genes (see methods, **Fig. 1B** and **Additional file 3: Table S3**). We used the CCLE_TIL10 and BPRNACan signature matrices with the EpiDISH method to estimate cell composition in different samples with gene expression datasets (**Additional file 2: Table S1**). We benchmarked our approaches on multiple datasets against MCP-counter, quanTIseq and a recently developed deconvolution method that exploits signatures derived from tumour sample scRNAseq datasets: CIBERSORTx (Newman et al., 2019) (**Additional file 2: Table S2**).

### Validation of the CCLE_TIL10 and BPRNACan signatures on tumor samples

After estimating performance of the signatures and methods on in-silico samples in which true cell proportions are known (**Additional File2: Supplementary Text S2, Figures S3a,b**), we investigated whether these deconvolution signature matrices would be able to estimate tumor purity from real biological samples, namely tissue samples from TCGA (**Additional File 2, Table S1**). We therefore analysed the results obtained with the 2 signature matrices (CCLE_TIL10 and BPRNACan) on TCGA-LUAD samples (**Additional File 2, Figure S4**). We first compared the estimation of cancer cell proportions in the samples to values estimated from 4 different methods (ABSOLUTE, ESTIMATE, LUMP and IHC, see methods). We found that the tumor purity estimates derived using BPRNACan were better than those derived from CCLE_TIL10 signature matrix on the TCGA-LUAD dataset, with correlations with the ESTIMATE method reaching Pearson’s R = 0.72 (**Additional File 2, Figure S4**).

As far as different immune cell types are concerned, we compared our deconvolved proportions using different methods and signatures to estimations of lymphocytes, macrophages and neutrophils based on H&E images (Saltz et al., 2018) (**Additional file 2: Table S1**). (**Fig. 2**). Generally, the best estimates for lymphocytes and macrophages over all cancer types were produced by the CIBERSORTx and BPRNACan signatures using either the CIBERSORTx method or EpiDISH. For neutrophils estimates, we observed the correlation with different signatures was very variable, probably due to the presence of neutrophils inside tumors of specific types having a specific phenotype not captured by all signatures. Overall, the best signature for neutrophils was the LM22 signature with our BPRNACan signatures capturing neutrophils specifically in Rectum Adenocarcinoma. quanTIseq estimates showed particularly low correlations with H&E estimates for neutrophils (**Fig. 2**).

**Figure 2.**
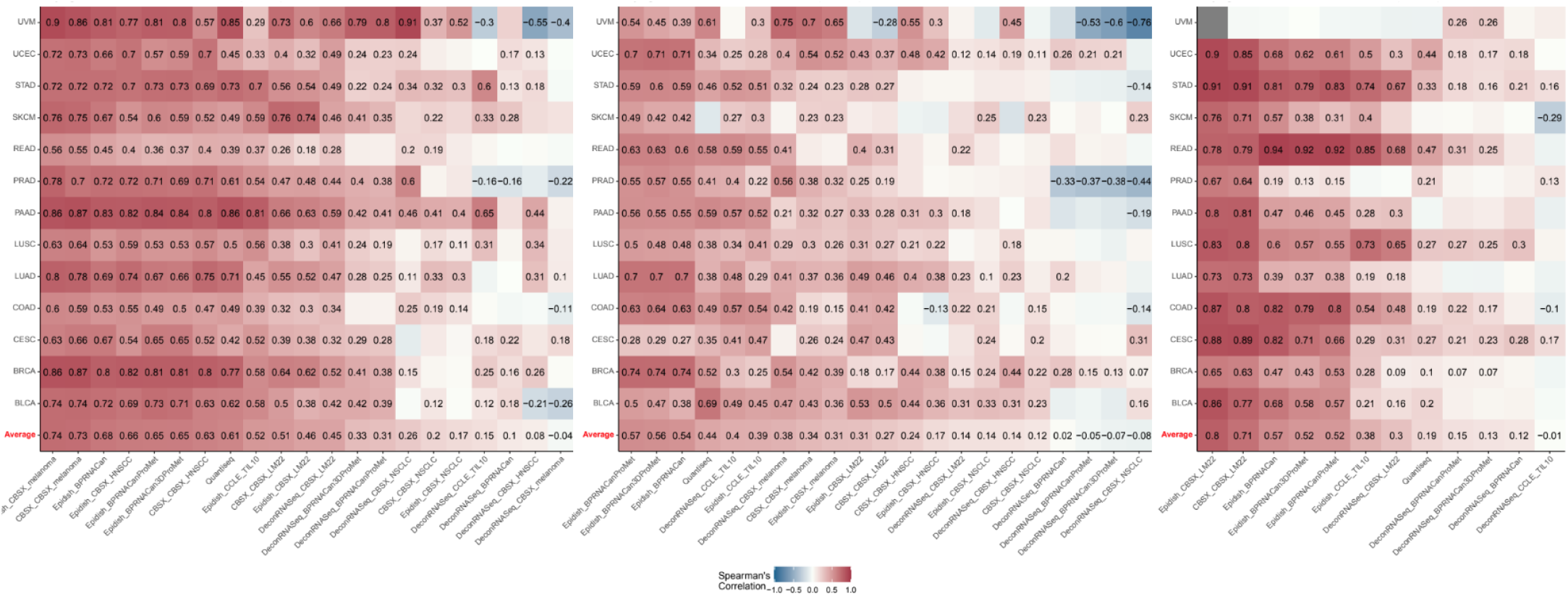
Comparison of different signatures for estimation of different cell types proportions across TCGA datasets. The heatmaps show Pearson correlation of deconvolved estimates and H&E imaging based estimates. Lymphocytes (left), Macrophages (center) and Neutrophils (right).

### A Novel DNA methylation-based signature matrix for immune and cancer cell deconvolution

Aiming to ultimately quantify the proportion of cancer and immune cells in a bulk sample from DNA methylation, we exploited the WGBS methylation datasets that were produced for bulk samples of purified cells as part of the Blueprint project (Farlik et al., 2016) to generate a signature matrix of 502 CpGs identifying 6 major immune cell types in blood (see **Additional File 1: Table B**). We named this signature matrix BPmet (**Additional File 2, methods**, **Additional file 3: Table S4**). To test this newly generated signature matrix, we performed deconvolution with the EpiDISH R package (Teschendorff et al., 2017), using our BPmet signature matrix, and choosing the RPC method as well as other available methods (**Additional file 2: Table S2**). We tested our BPmet new signature matrix on various datasets (**Additional file 2: Table S1**) identifying equal or better performance compared to other existing methods with a Pearson correlation between estimates and FACS measurements above 0.93 (**Additional file 2: Supplementary figure S5,6**).

Confident of the satisfying performance of the BPmet signature matrix in identifying cell type proportions in whole blood samples, we turned towards applying deconvolution to cancer tissues, extending the BPmet signature matrix to also estimate the proportion of cancer cells. We combined data from the immune samples that were used for the development of BPmet (**see methods**) with 17 normal and 34 cancer samples to derive a combined signature (see methods for details). This signature matrix, called BPmetCan (see methods, **Fig. 1b and Additional file 3: Table S5**), allowed us to estimate the proportion of cancer cells as well as of 10 immune cell types in bulk tumor samples.

To test the performance of BPmetCan in a realistic scenario, we used it to estimate the proportion of cancer cells in a subset of TCGA datasets with DNA methylation (15 cancer types, see **Additional file 2: Table S1**). Firstly, we retrieved publicly available estimates of the proportion of cancer cells in these samples (purity) produced using the ABSOLUTE, ESTIMATE, LUMP and IHC methods. When comparing purity estimates using BPMetCan with ABSOLUTE results, we obtained a Pearson correlation of R = 0.78 and P < 2.2e-16, proving that our signature matrix is able to identify the proportion of cancer cells in a tumor sample (**Additional file 2: Figure S7a**).

### Validation of the BPmetCan signature matrices on tumor samples

In order to assess our ability to determine the proportions of other immune cell types in the sample, the estimated fraction of immune cells using BPmetCan or MethylCIBERSORT signature matrix and the EpiDISH or CIBERSORT methods were compared to quantification of lymphocytes, macrophages and neutrophils estimated from H&E images also in the same samples from TCGA (Saltz et al., 2018). We observed that BPmetCan and MethylCIBERSORT have very similar performances, as measured by the correlation of deconvolved immune cell types compositions with H&E estimates (**Fig. 3)**. Investigating macrophages (Monocytes/Macrophages lineage) showed that the prediction of macrophage abundance derived from MethylCIBERSORT was more accurate than that derived using the BPmetCan signature matrix (R = 0.48 against R = 0.33) (**Additional File 2: Figure S7b**).

**Figure 3.**
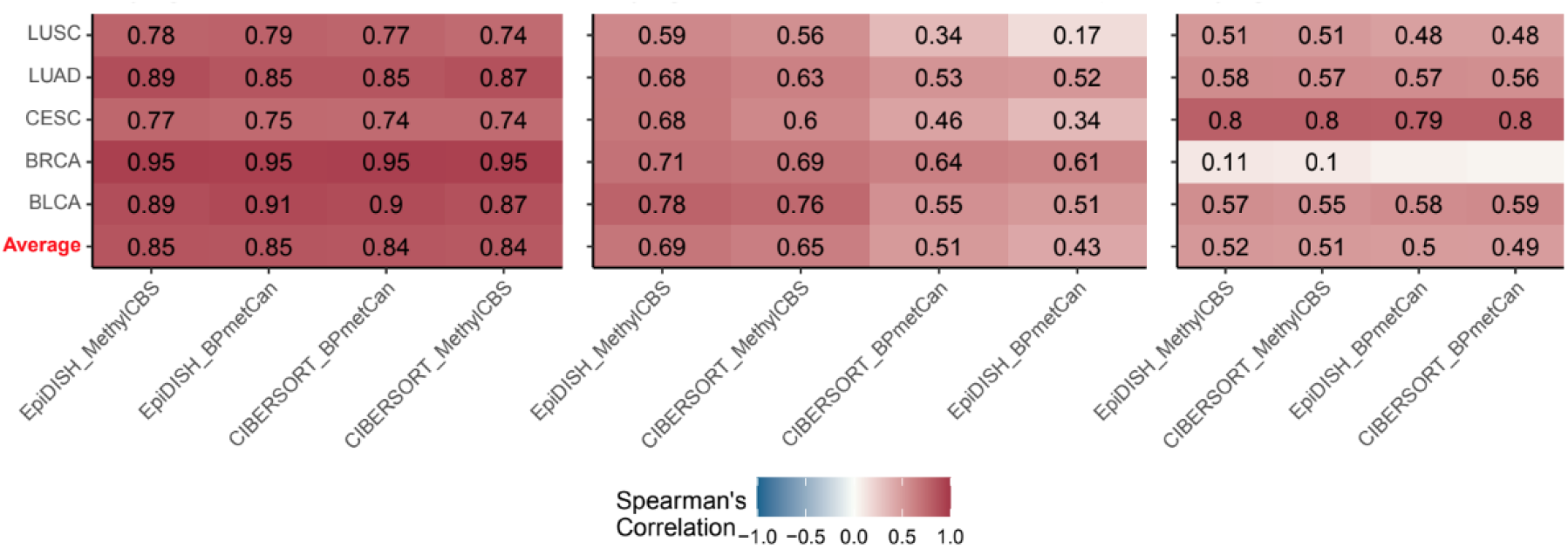
Comparison of different signatures and methods for estimation of different cell types proportions across TCGA datasets using DNA methylation. The heatmaps show Pearson correlation of deconvolved estimates and H&E imaging based estimates. Lymphocytes (left), Macrophages (center) and Neutrophils (right).

Nevertheless, it appears that the MethylCIBERSORT reference profiles do not capture all macrophages (some samples are on the x-axis indicating no estimated abundance while non-zero values are clearly detected by H&E images). This could be due to the classification of macrophages by the MethylCIBERSORT signature matrix that only includes monocytes (CD14+ cells), whereas our signature matrix can potentially classify monocytes, M0, M1 and M2 separately (all these types were merged for making this comparison).

### Gene expression and methylation based cell type signatures are significantly associated

As we have seen, expression and methylation data can contain complementary information that can be useful to deconvolve cell type proportions. Thinking about practical situations and in particular translational applications in the clinic, these two data types are rarely available for the same samples. At first, we thought of considering whether we could improve our gene expression based signatures by considering our knowledge of DNA methylation specificities of the different cell types that are reflected by the CpGs included in the DNA based signatures. We therefore investigated what could be the relation between the genes that are included in the BPRNACan signature matrix gene set (1403 *sig genes)* and the CpGs from the BPMetCan signature matrix (1896 *sig CpGs)* to assess whether they referred to the same or different genes. We calculated the number of *sig genes* that are associated with the *sig CpGs*. To associate CpGs to genes we first used Illumina annotation, which provides gene associations for each CpG on their array, based on genomic proximity between the gene and the CpG. Out of 169 genes associated with *sig CpGs*, 24 were included in the 1403 *sig genes* (Fisher’s exact test p-value=1e-5) (**Additional File 2: Table S3a**).

### Exploiting 3D chromatin interactions maps to combine methylation and expression based signature matrices

Inspired by the availability of 3D chromatin contact maps for haematopoietic cells produced by the Blueprint project (Javierre et al., 2016) and by known relations between 3D conformation, methylation and expression levels (Madrid-Mencía et al., 2020), we decided to re-evaluate CpGs annotation to genes and consider 3D chromatin contacts between these CpGs and gene promoters, as identified by Promoter-Capture Hi-C (PCHi-C). Briefly, this method involves a step of hybridization to a promoter library during a traditional Hi-C experiment, resulting in chromatin contact maps that contain either contacts amongst promoters or between a promoter and a regulatory region (denoted as OE for Other end or PIR for Promoter Interacting Region), where chromatin fragments have a median size of 5kb (Schoenfelder et al., 2015). The total PCHi-C network for haematopoietic cells consists of 249,511 chromatin fragments, of which 20,582 are promoter fragments (including 1127 fragments of *sig gene* promoters) and 228,929 other ends, (including 1131 fragments containing *sig CpGs*) (**Additional file 2: Table S4**).

We first mapped *sig CpGs* to their corresponding gene promoters, according to the definition of promoters used in the PCHi-C datasets. We found 645 genes that have *sig CpGs* in their promoter, of which 52 were already identified in *sig genes* and 314 were not in *sig genes* and had expression profiles in our reference datasets. Moreover, the comparison of the mapping between *sig CpGs* and genes harboring *sig CpGs* in their promoter shows that the PCHiC based annotation of CpGs to promoters reveals a higher overlap than Illumina annotation, Fisher’s exact test p-value = 7.34e-08 (**Additional File 2: Table S3b**), suggesting that associating CpGs to only the closest promoter could be limiting.

We reasoned that genes whose promoter methylation is cell-type specific should be important, even if they are not included in the gene expression signature. We therefore created an expanded gene expression deconvolution signature matrix by adding these 314 genes, that had *sig CpGs* in their promoters but were not *sig genes*, to the BPRNACan signature, leading to the BPRNACanProMet signature (**Figure 4, Additional file 3: Table S6**).

**Figure 4:**
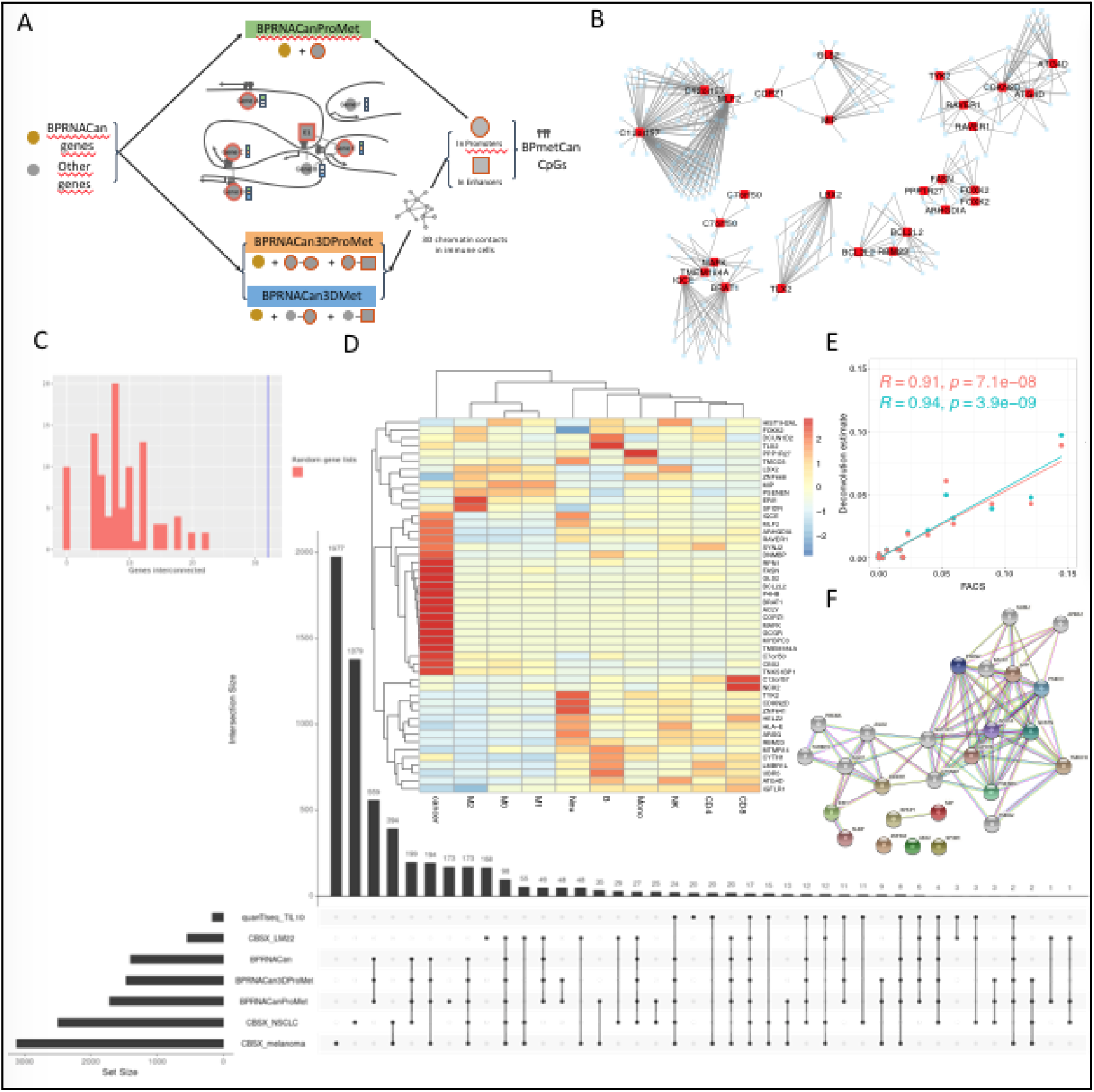
Strategy of integration of BPRNACan and BPmetCan signature matrices via 3D chromatin contact networks. (**a**) The BPRNACan signature is expanded including genes that have sig CpGs in their promoters (BPRNACanProMet), genes that have sig CpGs only in 3D regions in contact with their promoters (BPRNACan3DMet) and genes that have sig CpGs in both promoter and 3D contacting regions (BPRNACan3DProMet). (**b**) The expanded signatures and more particularly BPRNACan3DProMet is composed of 63 new genes some of whom are connected in a 3D chromatin contact map. C) The number of contacts amongst the 48 genes that are unique to our BPRNACan3DProMet and BPRNACanProMet signature compared to the number of contacts in random gene lists of equal size. D) UpsetR plot showing overlap between different gene signatures. Inset: heatmap of expression of those 48 genes investigated in C. E) Comparison of correlation between FACS estimates and the deconvolved proportion of macrophages using BPRNACan (red) and BPRNACan3DProMet (cyan). F) Functional interaction network of the 6 genes that are uniquely present in our BPRNACan3DProMet signature and discriminative of the M1 and M2 subtypes generated by String.

Given the importance of gene regulation by genomically distal regulatory elements that are brought into 3D contact with the promoter, having *sig CpGs* in both the promoter and a distal interacting fragment could increase the benefit of including the gene in the gene expression deconvolution signature. We therefore considered a more stringent expansion of our BPRNACan signature matrix to include only the genes that have *sig CpGs* both inside their promoter and in regions that contact their promoter in 3D (BPRNACan3DProMet, **Additional File 3: Table S7**).

The principle of gene inclusion in these 3 signatures is detailed in **Fig. 4a**, where we also show that genes newly included in our signature tend to connect in 3d (**Fig. 4b**) more than expected at random (**Fig. 4c**) and the expression of a subset of these genes found by studying overlaps between the different signatures (**Fig. 4d**) can clearly separate macrophage subtypes **(Fig. 4d inset)**.

We found the BPRNACanProMet and the BPRNACan3DProMet signature matrices to have better performance in estimating macrophage proportions compared to the original BPRNACan signature matrix, especially (**Fig. 4e, Fig. 2 center**). Intrigued by what could be the impact of considering these genes with altered epigenome in their promoter and distal interacting regions, we looked for genes that are exclusively present in the BPRNACan3DProMet signature. In the extra 63 genes that are added to the signature in this fashion, there are some genes involved in immune processes like HLA-E, CD4, P4HB (interacting w/ Th2 T Cells), TYK2, NCK2 (interleukin 23 signaling), or genes involved in hematopoiesis such as PTPN6, LMBR1L. Other genes are markers of cancer like ARHGDIA, CYTH1. Some of these genes present in the BPRNACan3DProMet signature were already present either in the BPRNA signature or in others that we have investigated in this paper. However 48 genes are uniquely present in the BBRNACan3DProMet signature (**Fig. 4f**). The biggest impact of the expansion of gene lists based on 3D contacts is on the macrophage signature, suggesting that potentially some of the genes captured by considering DNA methylation changes in their neighbourhood could distinguish tumour associated macrophages of different types.

The 6 macrophage specific genes, namely SPIDR, ERI1, PSENEN, MIP, ZNF668 and LBX2 were found to interact with NOTCH1(PSENEN) and DICER (ERI1) we thus investigated whether the NOTCH pathway and RNA silencing had been associated to specific macrophage subtypes. NOTCH has been associated with macrophage polarization and phagocytosis, while DICER conditional deletion has been recently implicated in loss of M2 polarization, via hyperactivation of IFNg. Interestingly these findings suggest that our signature could be specific to tumour protective M2-like macrophages, which play an important role in resistance to immunotherapy in patients.

### Combining deconvolved proportions based on gene expression and DNA methylation data

Since deconvolution can be performed using both expression and methylation data, we next worked on combining results using these two data types, assessing whether the performance of the combined analysis was better. This approach could be particularly useful in the cases for which both gene expression and DNA methylation data are available on the same samples or patients, where we can simply use a combined estimate for cell type abundance.

Exploiting the availability of both DNA methylation and gene expression data in TCGA, we evaluated the performance of our combined approach on these data. As shown in **Figure 5,** the correlation between deconvolved lymphocyte proportions and H&E estimations is much higher using DNA methylation than using RNAseq data (an average across cancer types of Pearson’s r = 0.86 vs 0.65), but it is still marginally improved when using an average between the two estimates (r=0.87). As far as macrophages are concerned, performances are less uniform across cancer types, but the estimations provided by deconvolution based on RNAseq, especially the BPRNACan3Dpromet signature, are better than those resulting from DNA methylation data in all cancer types but SKCM and are not improved by taking the average of methylation and expression based estimates. For neutrophils, the performance is again very variable across cancer types but overall it is improved in most cases when considering a combination of estimates based on DNA methylation and expression.

**Figure 5:**
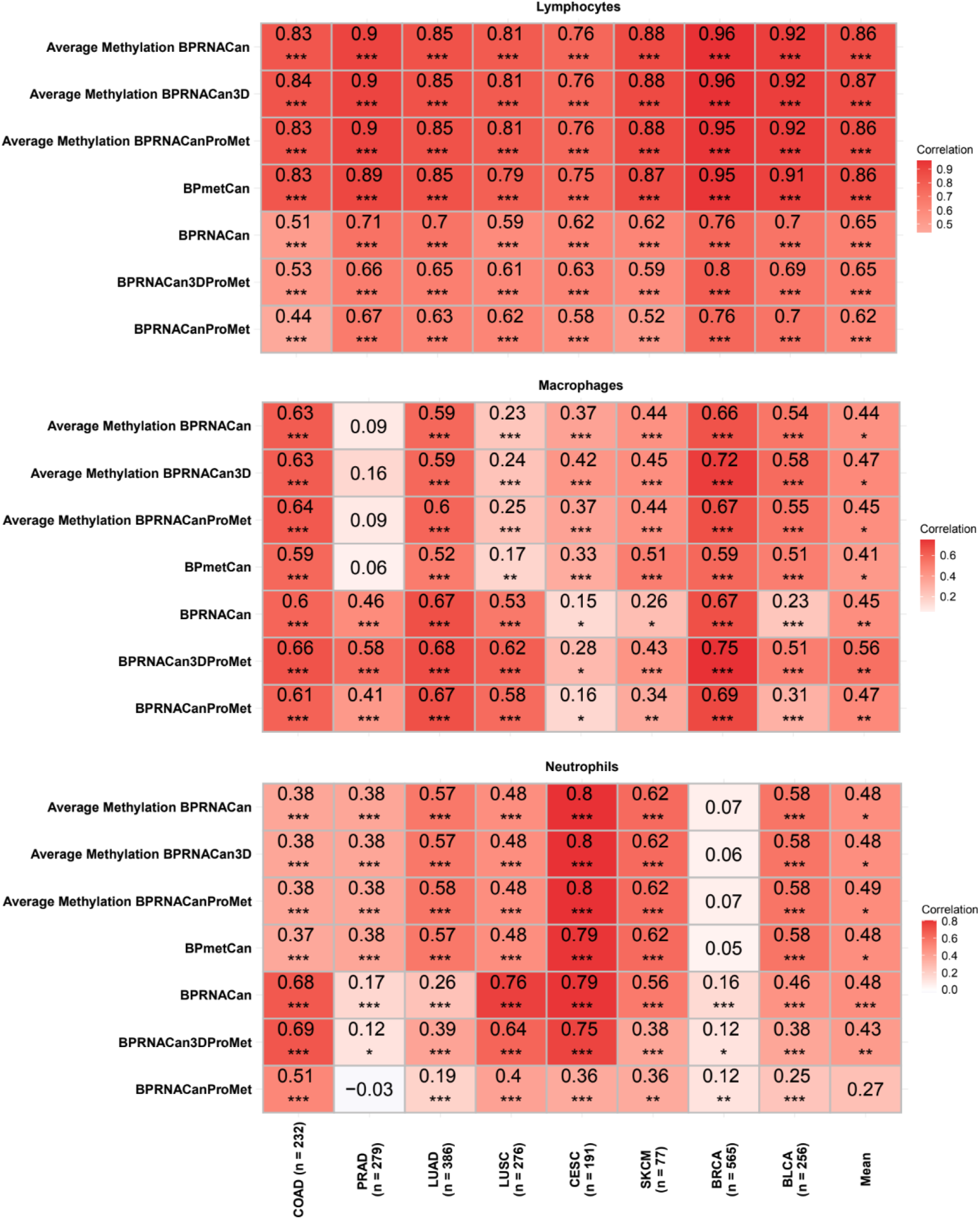
Performance on TCGA datasets of DNA methylation based, expression based and combined deconvolution. Correlation of estimates based on H&E staining is shown as cell colour, significance is indicated by asterisks.

### Benchmarking different methods and signatures in tumour samples with known composition

To overcome the limitation of using only rough estimates of cell types composition (namely identifying lymphocytes, macrophages and neutrophils ignoring all the important cellular subtypes), we turned towards datasets for which the composition was ascertained by either single cell sequencing of the tumour samples.

We examined an scRNAseq dataset including 19 primary melanoma non metastatic samples (Tirosh et al., 2016) and aggregated reads to produce an *in-silico* bulk sample for each patient to compare deconvolved proportions to the corresponding cell fractions estimated by counting single cells of each type from Racle et al. (Racle et al., 2017) (CD8 T cells, macrophages, B cells and NK cells). We ran deconvolution using all combinations of methods and signature matrices.

We observed that for three separate methods (EpiDISH, CIBERSORTx and DeconRNAseq) the signature derived from single-cell melanoma data performs the best on average across cell types. The BPRNACan signatures (specifically in order BPRNACan, BPRNACanProMet and BPRNACan3DProMet) are the next most performing ones. Interestingly, for almost all specific cell types the BPRNACan signatures perform the best, their correlations being lower than the CIBERSORTx signatures only for Macrophages and NK cells. Since macrophages are broadly defined including both M1 and M2 polarised cells, we could not test the ability of our signatures to distinguish these two important cell subtypes.

### Using deconvolution to predict response to immune checkpoint inhibitors

We have so far compared performances of different deconvolution methods and signatures based on cell type composition estimates provided by FACS, H&E image quantification of specific markers or single-cell data. All of these methods, however, carry their own biases in estimating composition. One of the main reasons that we are interested in quantifying cell types in the TME is to better understand and predict response to immune check-point inhibitors in a personalized medicine framework. This remains challenging and biomarkers of response are needed to achieve better outcomes in immunotherapy. We therefore decided to test the pertinence of our signature matrices to produce features for models that predict response to immunotherapy based on bulk transcriptomics (**Fig. 6a**).

**Figure 6:**
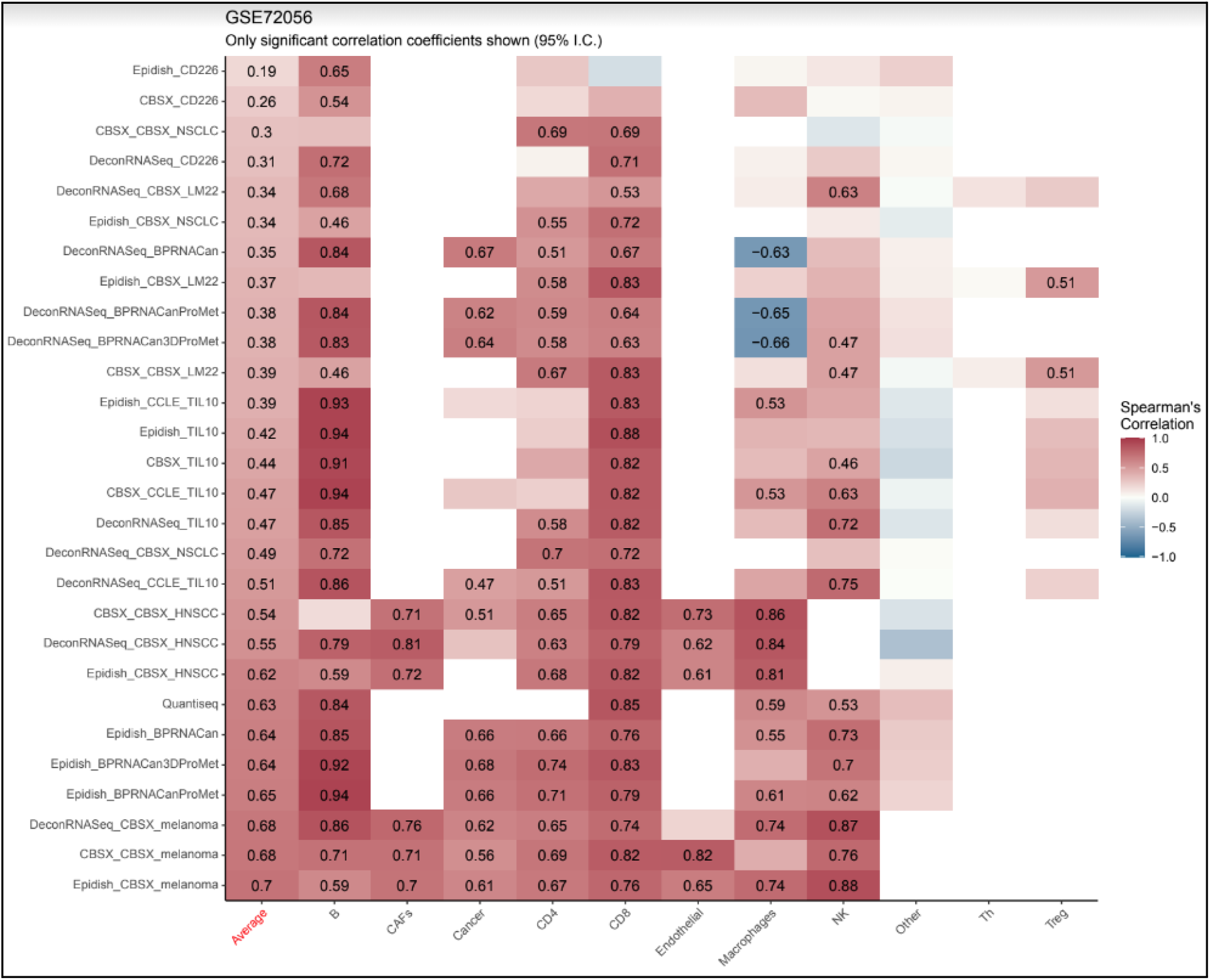
Performance of different methods and signature combinations on an RNAseq dataset from 19 melanoma samples for which cell type composition was estimated by scRNAseq performed on the same sample (Tirosh et al. 2016).

To this end, we made predictors of response to anti-PD1 antibodies using different deconvolution results as features and evaluated their performance on 3 public melanoma datasets (Gide et al., 2019; Hugo et al., 2016; Riaz et al., 2017) with response to anti-PD1 through ElasticNet penalized logistic regression (see **Methods** and **Fig. 7**).

**Figure 7:**
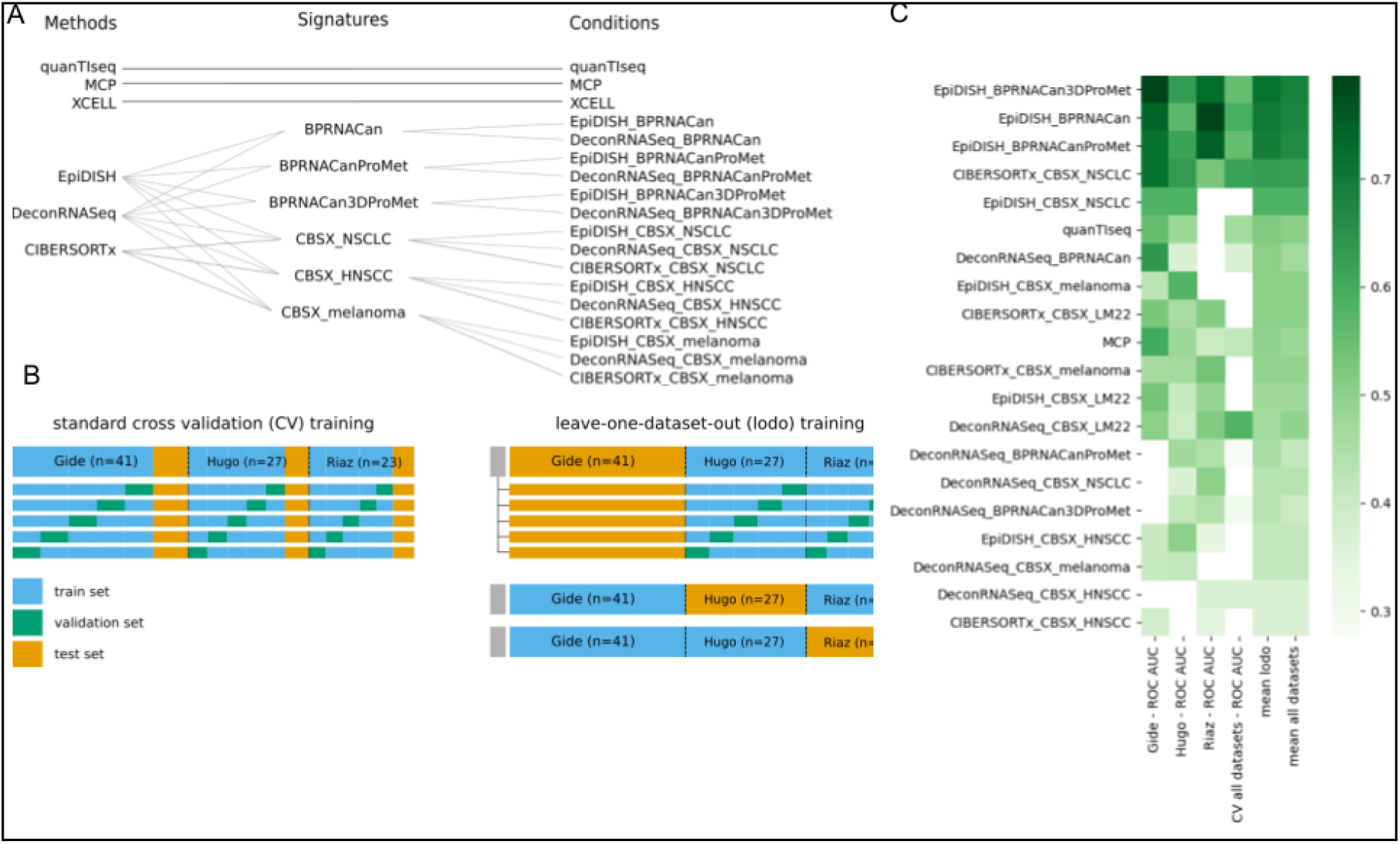
Evaluation of predictive power of deconvolution features. (**a**) Each combination of signature and deconvolution method was used alone to estimate cell types proportions for all samples. This data was used to train and test an elasticnet penalized logistic regression model (Friedmann et al. 2010) to predict response to immunotherapy. Performance of models can then be compared across signatures and training methods. (**b**) **Left:** The first training method is standard cross validation, during which a fourth of all samples is held-out as a test set. The remaining samples are used for training, with a 5-fold CV search for the hyperparameters that includes successive training sets (blue) and validation sets (green). **Right:** An alternative training method is leave-one-dataset-out (lodo), where one dataset is used for testing (orange) and the other ones are used for training (blue). During each training phase, hyperparameters for the l1 ratio and regularization strength were searched for by 5-fold cross validation (CV), where the training set is subdivided in 5 parts, each one of them being used as a validation set (green) while the other ones are used for training. At the end of the CV search the model is re-trained on all training samples, and tested on the test set. (**c**) Table of models’ performances by combinations of signatures and deconvolution methods and by training methods. The performance is assessed with the ROC AUC score computed on test sets, which is indicated by the name of the dataset for the lodo training. For the standard CV, a fifth of samples was held out as a test set.

Predictive models trained from cell type proportions produced by EpiDISH in combination with our signatures were the top performing models, followed closely by CIBERSORTx used in combination with the CBSX signature of NSCLC (**Fig 7c**). These models are also the top performing ones when considering AP and MCC as performance metrics (data not shown).

Our signature matrices in combination with EpiDISH or DeconRNASeq allowed us to train models with a performance that was better than random in a classification task (Receiving Operator Characteristic (ROC) Area Under the Curve (AUC) score above 0.5 on the standard Cross Validation (CV) training for 10 models among 12). On specific datasets unseen during training (see **Methods**) our signatures reached AUC=0.77, while training on all samples (standard CV), the BPRNACanProMet signature used by EpiDISH was the top performing model, with a ROC AUC score of 0.703 (**Fig. 7c**).

We then inspected the top 4 performing models’ coefficients, positive coefficients being associated to progressive disease. Models trained on cell type proportions estimated by our 3 signatures with EpiDISH have identical top 3 coefficients: M2 are strongly associated with progressive disease, whereas M0 and B cells are associated with response to immunotherapy. The remaining coefficients are smaller, and models agree on M1 being slightly associated with progressive disease, but the role of NK cells and neutrophils differs between models. The 4^th^ top performing model, based on features produced by CIBERSORTx with the CBSX NSCLC signature, shows CD4 cells strongly associated with response to immunotherapy, and monocytes associated to progressive disease. In this model, NK cells are also associated with progressive disease, which is in contradiction with previous models.

Looking at the coefficients in our regression models we can estimate which variables are associated with either progressive diseases or response to therapy. The coefficients of the model trained with the combination of the BPRNACanProMet signature and the EpiDISH method show that B cells (presumably indicative of the presence of tertiary lymphoid structures) and M0 macrophages proportions are associated with patient response, whereas M2 macrophages, CD4, CD8 and NK cells associated with progressive disease in this model. Proportions of cancer cells, M1 macrophages, monocytes and neutrophils were not significantly associated with either of the two outcomes. We can also consider the combination of the BPRNACan signature with deconRNASeq that always produced models performing better than a random predictor, and that is the second best when the Snyder dataset was used as a test set. The coefficients of this model indicate that B cells, M0 and M1 macrophages proportions are associated with response of patients, whereas CD4 and CD8 cells, Nk cells, monocytes and M2 macrophages are associated with progressive disease in this model. Like the previous model, proportions of cancer cells and neutrophils were not strongly associated with any of the two outcomes. Thus, the two models differ in the importance of M1 macrophages and monocytes for predicting response or progressive disease.

## Discussion

Numerous reference-based deconvolution methods using DNA methylation or gene expression can be used to estimate the proportion of cell types in bulk datasets from cell mixtures, such as EpiDISH, MethylCibersort (for DNAm) and CIBERSORT, MCP-counter, quanTIseq, DeconRNASeq (for GE). Moreover, there are methods that can accurately predict the proportion of cancer cells in tumor samples (purity). However, for application in immuno-oncology it can be important to estimate the proportions of cancer and specific immune cells at the same time.

Recently, Chakravarthy *et al*. (Chakravarthy et al., 2018) have reported that deconvolution methods based on gene expression or DNA methylation could be complementary to each other in cases where both data types are available. Despite this, they did not further explore if the reference signature matrices for each type of data can be related. Often in the clinic only gene expression data is available, so the integration of the two approaches is not common. Here, we present novel signature matrices for reference-based deconvolution:

**BPmetCan:** which is able to deconvolve the proportion of cancer and immune cell types from DNA methylation data (Illumina Arrays or WGBS), based on WGBS signature matrices, which we validate against MethylCIBERSORT, EpiDISH and various methods to estimate tumor purity in blood and especially in tumor samples.
**CCLE_TIL10:** which combines the TIL10 signature matrix (Finotello et al. 2019) for immune cell types with a new list of genes identified to be cancer-cell specific using data from GTEx and CCLE, which displays excellent performance on in-vitro and in-silico mixtures of cancer cell lines and immune cells.
**BPRNACan:** which combines our Blueprint derived immune cell signature matrix (BPRNA) with genes that are discriminant of cancer tissues compared to normal and outperforms or equals quanTIseq and MCP-Counter on cancer samples for many of the cell types.
**BPRNACanProMet:** which is an enhancement of the BPRNACan signature matrix by adding genes that have a *sig CpG* contained in their promoter, with demonstrated improvement in performance on many datasets
**BPRNACan3DProMet:** which is an enhancement of the BPRNACan signature matrix by adding genes that have a *sig CpG* in their promoter and whose promoters also have a 3D contact with a fragment containing a *sig CpG*.

We performed extensive validation using previous studies, such as whole blood mixtures, solid tumor-TCGA, PBMC, multiple myeloma patient bone marrow samples and melanoma non metastatic and metastatic samples. We compared the available DNAm or GE signature matrices to demonstrate that our novel signature matrices could faithfully estimate the fraction of cancer and specific immune cell compositions from DNA methylation and bulk gene expression data. Our signature matrices can be applied to solid tumors, as confirmed by the validations presented above, but they are likely to have limited use for hematological malignancies, in which the presence of cancerous immune cells could confound the estimations, and for which more targeted signature matrices should be developed.

We also showed that our DNAm signature matrix BPmet is more robust than others for estimating the proportions of cell types in Whole Blood samples. Moreover, our new gene expression signature matrix BPRNACan displayed higher accuracy on the predicted cell fractions on in-vivo cancer samples. Additionally, application of our GE signature matrices to publicly available data using our predicting model in this study revealed several important biological insights on predicting response to immunotherapy.

Despite the overall accuracy of our signature matrices, we found that the performance in the estimation for some cell types is lower than what we expected, probably due to the number or condition of the cell types used for establishing the reference profiles. For instance, we observed that the correlation of NK cells was lower than expected using both BPmet and BPRNA. This may be explained as we had only n = 2 samples to create the NK reference expression profile. Moreover, we observed a very low performance in predicting M2 macrophages in the *in-silico* tumor samples using BPRNACan. This may be explained by the fact that this M2 state in-vitro is artificially induced by cytokines to mimic the context of the TME, thereby the GE reference profiles corresponding to M2 cells might not capture the M2s that are found, albeit at low frequency (Shen-Orr and Gaujoux 2013), in artificial *in-silico* mixtures of purified cells. As opposed to NK and M2 cells, CD8 T cells and Macrophages were better predicted using GE signature matrices on cancer samples. This could be explained by the differences in cell proportions and cell states between the tumor microenvironment and circulating blood. For example, only a few activated CD8 T cells and no macrophages can be found in circulating blood, but their detection in tumors is key.

We also found a low performance in detecting neutrophils according to proportions derived by H&E (**Fig. 2**). This may be explained by the fact that the reference profiles of neutrophils used to build the signature matrix are unlikely to capture all the phenotypes that neutrophils can display, especially inside tumors. Like macrophages, Tumor Associated Neutrophils (TANs) can display at least two different phenotypes - one characterised by proinflammatory programs and antitumorigenic functions and the second characterised by a protumorigenic activity (Shaul and Fridlender 2019; Teijeira et al. 2020). Indeed neutrophils’ gene expression and methylation were found to be extremely variable across individuals (Ecker et al. 2017), time of the day, as well as across the different parts of the body in which they are found, highlighting their extreme plasticity (Giese, Hind, and Huttenlocher 2019; Sagiv et al. 2015).

Another important novel insight of our study is that the GE signature matrix can be improved by incorporating specific genes that are associated with CpGs included in the DNA methylation signature matrix. This can be done by either identifying genes that harbor signature matrix CpGs (*sig CpGs*) in their promoter, or those genes whose promoters also have a 3D contact with a fragment containing a *sig CpG*. We are thus able to expand the list of genes to be used in performing RNAseq-based deconvolution using information gathered from the DNA methylation signature matrix we have generated (**Additional file 2: Table S2**). Importantly, this expanded GE signature matrix can be applied to improve deconvolution of samples for which only RNAseq data is available (**Figure 5**).

Despite our DNAm and RNAseq reference-based signature matrices displaying comparable to or better performance than existing methods in whole blood or cancer samples, several issues will require further investigation. We are limited by the number of samples available for specific cell types (such as NK cells) and by the fact that these profiles are generated from purified cells that are isolated from their natural environment. This is especially true for cells that acquire specific phenotypes in a TME context, such as TAMs and TANs. The availability of single-cell RNAseq datasets from cancer samples, especially from technologies that also provide protein marker quantification such as CITEseq, will greatly improve our chances of generating relevant signature matrices for any cell type of interest. For this reason we provide code for the generation of new reference signature matrices in our openly available repository. As a step in this direction, we have included the CIBERSORTx method using 3 scRNAseq derived signatures in our benchmark, despite the difficulty of integrating it in our openly available pipeline.

Importantly, one of the main applications of deconvolution in immuno-oncology will be the prediction of response to immunotherapy, which can be made based on the inferred cell type proportions. We therefore benchmarked our new signature matrices and methods by evaluating their accuracy in predicting response to anti-PD1 agents in 3 public datasets. This type of exercise is aimed at identifying which signature matrices and methods uncover the presence of specific cell subtypes that can impact immune checkpoint blocker response. These important subtypes might not correspond easily to literature definitions or FACS derived populations defined by specific markers. M2 macrophages are known to impair response to immune checkpoint blockade (Ceci et al., 2020) and we confirm their association with negative prognosis despite the discrepancy between our quantification of M2s and FACS estimates. This could be due to our signature capturing specific phenotypes that are not entirely determined by traditional cell surface markers used in FACS. Interestingly, B cells and M1 macrophages have recently been proposed as potential predictors of response to immunotherapies (Cabrita et al., 2020; Helmink et al., 2020; Petitprez et al., 2020; Zeng et al., 2020). To our knowledge, the proportion of M0 (naïve) macrophages has not been reported as a predictor of response to immunotherapies, but both of our models suggest it is a potentially relevant subtype in this respect associated with response. For NK cells and neutrophils we do not reach a consensus amongst our different models. One of the top performing models using scRNAseq based signatures with the CIBERSORTx method suggests CD4 cells as associated with response, while monocytes and strangely NK cells would promote disease progression. These results have to be considered with caution, as response to immunotherapy was shown to depend on the proportion of sub-cell types like memory, effector, and senescent phenotypes or defined by the presence or absence of several proteins like PD-1, PD-L1, CTLA-4, LAG-3 or TIGIT on these cells’ surfaces, which are not captured by our signatures.

In summary, we have shown the potential of our gene expression signature matrices to estimate the presence of immune populations that can be predictive of the response to checkpoint blockade, bringing us closer to personalized approaches and revealing resistance mechanisms.

## Conclusion

We have presented and thoroughly validated novel deconvolution approaches using DNA methylation or gene expression data or a combination of the two. Our signatures show comparable performances in estimating the proportion of cell types from blood and tumor samples according to multiple benchmark datasets. Simultaneously, we have also shown that our signature matrices are more robust to estimate cell fractions compared to the other available signature matrices. We also showed how expanding a gene expression signature matrix using information regarding methylation patterns and 3D chromatin contacts between promoters and regulatory elements we can improve our signatures performance, especially for the detection of tumour associated macrophages. This improved macrophage signature has predictive power for estimating response to anti-PD1 agents in 3 publicly available melanoma datasets. We make the signature matrices and all the code available to the community through a user-friendly snakemake pipeline that can use any given reference signature matrix to apply a variety of reference-based deconvolution methods. Additionally, the code to generate a signature matrix following the method used in this paper is also available, as well as a script to generate all the figures.

Finally, we provide a user-friendly pipeline to apply our approach and make the code available to the research community, to promote the development of new, more specific deconvolution signatures (https://github.com/VeraPancaldiLab/GEMDeCan). Despite the increasing availability of single cell RNAseq data, we propose that deconvolution of cell types and subtypes from bulk transcriptomics will be a valid strategy to target immunotherapies to specific patients in a clinically relevant context and at an affordable cost.

## Abbreviations

BRCA: Breast invasive carcinoma
CCLE: Cancer Cell Line Encyclopedia
DHS: DNAse hypersensitivity sites
DNAm: DNA methylation
FACS: fluorescence-activated cell sorting
FPKM: Fragments Per Kilobase of transcript per Million
GE: gene expression
GEO: Gene expression omnibus
H&E: Hematoxylin and eosin
IHC: Immunohistochemistry
LUAD: Lung adenocarcinoma
M: macrophages
M1: Classically activated macrophages
M2: Alternatively activated macrophages
Mono: Monocytes
Neu: Neutrophils
NK: Natural killer cells
PBMC: Peripheral blood mononuclear cells
PCHi-C: Promoter-Capture Hi-C
R: Pearson’s correlation
RPC: robust partial correlation
TANs: tumor Associated Neutrophils
TAMs: tumor Associated Macrophages
TCGA: The Cancer Genome Atlas
TME: The tumor microenvironment
TPM: Transcripts per millions
Treg: Regulatory T cells
WB: whole blood
WGBS: whole-genome bisulfite sequencing

## Acknowledgements

We thank members of the Pancaldi lab for critical reading of the manuscript and Sarah Djebali for help with testing the pipeline.

## Availability of data and materials

The published data used is indicated in Methods. The pipeline to generate all examples is available from GitHub: https://github.com/VeraPancaldiLab/GEMDeCan and we provide an R notebook to reproduce all figures in the paper.

## Author information

**Centre de Recherches en Cancérologie de Toulouse (CRCT), INSERM U1037, Toulouse, France**

Ting Xie, Julien Pernet, Nina Verstraete, Miguel Madrid Mencía, Alexis Hucteau, Alexis Coullomb, Vera Pancaldi

**Université Paul Sabatier III, Toulouse 31400, Toulouse, France**

Ting Xie, Julien Pernet, Nina Verstraete, Miguel Madrid Mencía, Alexis Hucteau, Alexis Coullomb, Vera Pancaldi

**Translational Medicine, Institut de Recherche Pierre Fabre, Toulouse, France**

Mei-Shiue Kuo, Olivier Delfour, Francisco Cruzalegui

**Barcelona Supercomputing Center, Barcelona, 08034, Spain**

Miguel Madrid Mencía, Vera Pancaldi

## Corresponding author

Correspondence to: <u>Vera Pancaldi</u>

## Ethics approval and consent to participate

Not applicable.

## Consent to publication

Not Applicable.

## Additional files

**Additional file 1:**

List of datasets used in this study for DNA methylation and gene expression reference database construction. **TableA: BP RNAseq datasets**, **Table B: BP WGBS datasets, Table C: normal/cancer/blood methylation datasets.**

**Additional file 2: Supplementary Text and figures.**

**Additional file 3:** Seven novel deconvolution signature matrices BPmet, BPmetCan,

BPRNA, BPRNACan, CCLE_TIL10, BPRNACanProMet, BPRNACan3DProMet.

## References

Anderson, K. G., Stromnes, I. M., & Greenberg, P. D. (2017). Obstacles Posed by the Tumor Microenvironment to T cell Activity: A Case for Synergistic Therapies. Cancer Cell, 31(3), 311–325. https://doi.org/10.1016/j.ccell.2017.02.008

Aran, D., Sirota, M., & Butte, A. J. (2015). Systematic pan-cancer analysis of tumour purity. Nature Communications, 6, 8971. https://doi.org/10.1038/ncomms9971

Becht, E., Giraldo, N. A., Lacroix, L., Buttard, B., Elarouci, N., Petitprez, F., Selves, J., Laurent-Puig, P., Sautès-Fridman, C., Fridman, W. H., & de Reyniès, A. (2016). Estimating the population abundance of tissue-infiltrating immune and stromal cell populations using gene expression. Genome Biology, 17(1), 218. https://doi.org/10.1186/s13059-016-1070-5

Binnewies, M., Roberts, E. W., Kersten, K., Chan, V., Fearon, D. F., Merad, M., Coussens, L. M., Gabrilovich, D. I., Ostrand-Rosenberg, S., Hedrick, C. C., Vonderheide, R. H., Pittet, M. J., Jain, R. K., Zou, W., Howcroft, T. K., Woodhouse, E. C., Weinberg, R. A., & Krummel, M. F. (2018). Understanding the tumor immune microenvironment (TIME) for effective therapy. Nature Medicine, 24(5), 541–550. https://doi.org/10.1038/s41591-018-0014-x

Brinkman, A. B., Nik-Zainal, S., Simmer, F., Rodríguez-González, F. G., Smid, M., Alexandrov, L. B., Butler, A., Martin, S., Davies, H., Glodzik, D., Zou, X., Ramakrishna, M., Staaf, J., Ringnér, M., Sieuwerts, A., Ferrari, A., Morganella, S., Fleischer, T., Kristensen, V., … Stunnenberg, H. G. (2019). Partially methylated domains are hypervariable in breast cancer and fuel widespread CpG island hypermethylation. Nature Communications, 10(1), 1749. https://doi.org/10.1038/s41467-019-09828-0

Cabrita, R., Lauss, M., Sanna, A., Donia, M., Skaarup Larsen, M., Mitra, S., Johansson, I., Phung, B., Harbst, K., Vallon-Christersson, J., van Schoiack, A., Lövgren, K., Warren, S., Jirström, K., Olsson, H., Pietras, K., Ingvar, C., Isaksson, K., Schadendorf, D., … Jönsson, G. (2020). Tertiary lymphoid structures improve immunotherapy and survival in melanoma. Nature, 577(7791), 561–565. https://doi.org/10.1038/s41586-019-1914-8

Cairns, J., Freire-Pritchett, P., Wingett, S. W., Várnai, C., Dimond, A., Plagnol, V., Zerbino, D., Schoenfelder, S., Javierre, B.-M., Osborne, C., Fraser, P., & Spivakov, M. (2016). CHiCAGO: Robust detection of DNA looping interactions in Capture Hi-C data. Genome Biology, 17(1), 127. https://doi.org/10.1186/s13059-016-0992-2

Carter, S. L., Cibulskis, K., Helman, E., McKenna, A., Shen, H., Zack, T., Laird, P. W., Onofrio, R. C., Winckler, W., Weir, B. A., Beroukhim, R., Pellman, D., Levine, D. A., Lander, E. S., Meyerson, M., & Getz, G. (2012). Absolute quantification of somatic DNA alterations in human cancer. Nature Biotechnology, 30(5), 413–421. https://doi.org/10.1038/nbt.2203

Cavalli, G., & Heard, E. (2019). Advances in epigenetics link genetics to the environment and disease. Nature, 571(7766), 489–499. https://doi.org/10.1038/s41586-019-1411-0

Ceci, C., Atzori, M. G., Lacal, P. M., & Graziani, G. (2020). Targeting Tumor-Associated Macrophages to Increase the Efficacy of Immune Checkpoint Inhibitors: A Glimpse into Novel Therapeutic Approaches for Metastatic Melanoma. Cancers, 12(11), E3401. https://doi.org/10.3390/cancers12113401

Chakravarthy, A., Furness, A., Joshi, K., Ghorani, E., Ford, K., Ward, M. J., King, E. V., Lechner, M., Marafioti, T., Quezada, S. A., Thomas, G. J., Feber, A., & Fenton, T. R. (2018). Pan-cancer deconvolution of tumour composition using DNA methylation. Nature Communications, 9(1), 3220. https://doi.org/10.1038/s41467-018-05570-1

DeBerardinis, R. J. (2020). Tumor Microenvironment, Metabolism, and Immunotherapy. New England Journal of Medicine, 382(9), 869–871. https://doi.org/10.1056/NEJMcibr1914890

Engblom, C., Pfirschke, C., & Pittet, M. J. (2016). The role of myeloid cells in cancer therapies. Nature Reviews. Cancer, 16(7), 447–462. https://doi.org/10.1038/nrc.2016.54

Farha, M., Jairath, N. K., Lawrence, T. S., & El Naqa, I. (2020). Characterization of the Tumor Immune Microenvironment Identifies M0 Macrophage-Enriched Cluster as a Poor Prognostic Factor in Hepatocellular Carcinoma. JCO Clinical Cancer Informatics, 4, 1002–1013. https://doi.org/10.1200/CCI.20.00077

Farlik, M., Halbritter, F., Müller, F., Choudry, F. A., Ebert, P., Klughammer, J., Farrow, S., Santoro, A., Ciaurro, V., Mathur, A., Uppal, R., Stunnenberg, H. G., Ouwehand, W. H., Laurenti, E., Lengauer, T., Frontini, M., & Bock, C. (2016). DNA Methylation Dynamics of Human Hematopoietic Stem Cell Differentiation. Cell Stem Cell, 19(6), 808–822. https://doi.org/10.1016/j.stem.2016.10.019

Finotello, F., Mayer, C., Plattner, C., Laschober, G., Rieder, D., Hackl, H., Krogsdam, A., Loncova, Z., Posch, W., Wilflingseder, D., Sopper, S., Ijsselsteijn, M., Brouwer, T. P., Johnson, D., Xu, Y., Wang, Y., Sanders, M. E., Estrada, M. V., Ericsson-Gonzalez, P., … Trajanoski, Z. (2019). Molecular and pharmacological modulators of the tumor immune contexture revealed by deconvolution of RNA-seq data. Genome Medicine, 11(1), 34. https://doi.org/10.1186/s13073-019-0638-6

Finotello, F., Rieder, D., Hackl, H., & Trajanoski, Z. (2019). Next-generation computational tools for interrogating cancer immunity. Nature Reviews. Genetics, 20(12), 724–746. https://doi.org/10.1038/s41576-019-0166-7

Friedman, J., Hastie, T., & Tibshirani, R. (2010). Regularization Paths for Generalized Linear Models via Coordinate Descent. Journal of Statistical Software, 33(1), 1–22.

Ghandi, M., Huang, F. W., Jané-Valbuena, J., Kryukov, G. V., Lo, C. C., McDonald, E. R., Barretina, J., Gelfand, E. T., Bielski, C. M., Li, H., Hu, K., Andreev-Drakhlin, A. Y., Kim, J., Hess, J. M., Haas, B. J., Aguet, F., Weir, B. A., Rothberg, M. V., Paolella, B. R., … Sellers, W. R. (2019). Next-generation characterization of the Cancer Cell Line Encyclopedia. Nature, 569(7757), 503–508. https://doi.org/10.1038/s41586-019-1186-3

Gide, T. N., Quek, C., Menzies, A. M., Tasker, A. T., Shang, P., Holst, J., Madore, J., Lim, S. Y., Velickovic, R., Wongchenko, M., Yan, Y., Lo, S., Carlino, M. S., Guminski, A., Saw, R. P. M., Pang, A., McGuire, H. M., Palendira, U., Thompson, J. F., … Wilmott, J. S. (2019). Distinct Immune Cell Populations Define Response to Anti-PD-1 Monotherapy and Anti-PD-1/Anti-CTLA-4 Combined Therapy. Cancer Cell, 35(2), 238–255.e6. https://doi.org/10.1016/j.ccell.2019.01.003

Gong, T., & Szustakowski, J. D. (2013). DeconRNASeq: A statistical framework for deconvolution of heterogeneous tissue samples based on mRNA-Seq data. Bioinformatics (Oxford, England), 29(8), 1083–1085. https://doi.org/10.1093/bioinformatics/btt090

GTEx Consortium. (2013). The Genotype-Tissue Expression (GTEx) project. Nature Genetics, 45(6), 580–585. https://doi.org/10.1038/ng.2653

Hastie, T., Tibshirani, R., Sherlock, G., Eisen, M., Brown, P., & Botstein, D. (2001). Imputing Missing Data for Gene Expression Arrays. Technical Report, Stanford Statistics Department, 1.

Helmink, B. A., Reddy, S. M., Gao, J., Zhang, S., Basar, R., Thakur, R., Yizhak, K., Sade-Feldman, M., Blando, J., Han, G., Gopalakrishnan, V., Xi, Y., Zhao, H., Amaria, R. N., Tawbi, H. A., Cogdill, A. P., Liu, W., LeBleu, V. S., Kugeratski, F. G., … Wargo, J. A. (2020). B cells and tertiary lymphoid structures promote immunotherapy response. Nature, 577(7791), 549–555. https://doi.org/10.1038/s41586-019-1922-8

Houseman, E. A., Accomando, W. P., Koestler, D. C., Christensen, B. C., Marsit, C. J., Nelson, H. H., Wiencke, J. K., & Kelsey, K. T. (2012). DNA methylation arrays as surrogate measures of cell mixture distribution. BMC Bioinformatics, 13(1), 86. https://doi.org/10.1186/1471-2105-13-86

Houseman, E. A., Molitor, J., & Marsit, C. J. (2014). Reference-free cell mixture adjustments in analysis of DNA methylation data. Bioinformatics (Oxford, England), 30(10), 1431–1439. https://doi.org/10.1093/bioinformatics/btu029

Hugo, W., Zaretsky, J. M., Sun, L., Song, C., Moreno, B. H., Hu-Lieskovan, S., Berent-Maoz, B., Pang, J., Chmielowski, B., Cherry, G., Seja, E., Lomeli, S., Kong, X., Kelley, M. C., Sosman, J. A., Johnson, D. B., Ribas, A., & Lo, R. S. (2016). Genomic and Transcriptomic Features of Response to Anti-PD-1 Therapy in Metastatic Melanoma. Cell, 165(1), 35–44. https://doi.org/10.1016/j.cell.2016.02.065

Javierre, B. M., Burren, O. S., Wilder, S. P., Kreuzhuber, R., Hill, S. M., Sewitz, S., Cairns, J., Wingett, S. W., Várnai, C., Thiecke, M. J., Burden, F., Farrow, S., Cutler, A. J., Rehnström, K., Downes, K., Grassi, L., Kostadima, M., Freire-Pritchett, P., Wang, F., … Fraser, P. (2016). Lineage-Specific Genome Architecture Links Enhancers and Non-coding Disease Variants to Target Gene Promoters. Cell, 167(5), 1369–1384.e19. https://doi.org/10.1016/j.cell.2016.09.037

Koestler, D. C., Jones, M. J., Usset, J., Christensen, B. C., Butler, R. A., Kobor, M. S., Wiencke, J. K., & Kelsey, K. T. (2016). Improving cell mixture deconvolution by identifying optimal DNA methylation libraries (IDOL). BMC Bioinformatics, 17, 120. https://doi.org/10.1186/s12859-016-0943-7

Köster, J., & Rahmann, S. (2018). Snakemake-a scalable bioinformatics workflow engine. Bioinformatics (Oxford, England), 34(20), 3600. https://doi.org/10.1093/bioinformatics/bty350

Lawrence, T., & Natoli, G. (2011). Transcriptional regulation of macrophage polarization: Enabling diversity with identity. Nature Reviews. Immunology, 11(11), 750–761. https://doi.org/10.1038/nri3088

Madrid-Mencía, M., Raineri, E., Cao, T. B. N., & Pancaldi, V. (2020). Using GARDEN-NET and ChAseR to explore human haematopoietic 3D chromatin interaction networks. Nucleic Acids Research, 48(8), 4066–4080. https://doi.org/10.1093/nar/gkaa159

Mantovani, A., Marchesi, F., Malesci, A., Laghi, L., & Allavena, P. (2017). Tumour-associated macrophages as treatment targets in oncology. Nature Reviews. Clinical Oncology, 14(7), 399–416. https://doi.org/10.1038/nrclinonc.2016.217

Monaco, G., Lee, B., Xu, W., Mustafah, S., Hwang, Y. Y., Carré, C., Burdin, N., Visan, L., Ceccarelli, M., Poidinger, M., Zippelius, A., Pedro de Magalhães, J., & Larbi, A. (2019). RNA-Seq Signatures Normalized by mRNA Abundance Allow Absolute Deconvolution of Human Immune Cell Types. Cell Reports, 26(6), 1627–1640.e7. https://doi.org/10.1016/j.celrep.2019.01.041

Murray, P. J. (2017). Macrophage Polarization. Annual Review of Physiology, 79, 541–566. https://doi.org/10.1146/annurev-physiol-022516-034339

Nakamura, K., Kassem, S., Cleynen, A., Chrétien, M.-L., Guillerey, C., Putz, E. M., Bald, T., Förster, I., Vuckovic, S., Hill, G. R., Masters, S. L., Chesi, M., Bergsagel, P. L., Avet-Loiseau, H., Martinet, L., & Smyth, M. J. (2018). Dysregulated IL-18 Is a Key Driver of Immunosuppression and a Possible Therapeutic Target in the Multiple Myeloma Microenvironment. Cancer Cell, 33(4), 634–648.e5. https://doi.org/10.1016/j.ccell.2018.02.007

Newman, A. M., Steen, C. B., Liu, C. L., Gentles, A. J., Chaudhuri, A. A., Scherer, F., Khodadoust, M. S., Esfahani, M. S., Luca, B. A., Steiner, D., Diehn, M., & Alizadeh, A. A. (2019). Determining cell type abundance and expression from bulk tissues with digital cytometry. Nature Biotechnology, 37(7), Article 7. https://doi.org/10.1038/s41587-019-0114-2

O’Donnell, J. S., Teng, M. W. L., & Smyth, M. J. (2019). Cancer immunoediting and resistance to T cell-based immunotherapy. Nature Reviews. Clinical Oncology, 16(3), 151–167. https://doi.org/10.1038/s41571-018-0142-8

Pathria, P., Louis, T. L., & Varner, J. A. (2019). Targeting Tumor-Associated Macrophages in Cancer. Trends in Immunology, 40(4), 310–327. https://doi.org/10.1016/j.it.2019.02.003

Petitprez, F., de Reyniès, A., Keung, E. Z., Chen, T. W.-W., Sun, C.-M., Calderaro, J., Jeng, Y.-M., Hsiao, L.-P., Lacroix, L., Bougoüin, A., Moreira, M., Lacroix, G., Natario, I., Adam, J., Lucchesi, C., Laizet, Y. H., Toulmonde, M., Burgess, M. A., Bolejack, V.,. Fridman, W. H. (2020). B cells are associated with survival and immunotherapy response in sarcoma. Nature, 577(7791), 556–560. https://doi.org/10.1038/s41586-019-1906-8

Racle, J., de Jonge, K., Baumgaertner, P., Speiser, D. E., & Gfeller, D. (2017). Simultaneous enumeration of cancer and immune cell types from bulk tumor gene expression data. ELife, 6, e26476. https://doi.org/10.7554/eLife.26476

Riaz, N., Havel, J. J., Makarov, V., Desrichard, A., Urba, W. J., Sims, J. S., Hodi, F. S., Martín-Algarra, S., Mandal, R., Sharfman, W. H., Bhatia, S., Hwu, W.-J., Gajewski, T. F., Slingluff, C. L., Chowell, D., Kendall, S. M., Chang, H., Shah, R., Kuo, F., … Chan, T. A. (2017). Tumor and Microenvironment Evolution during Immunotherapy with Nivolumab. Cell, 171(4), 934–949.e16. https://doi.org/10.1016/j.cell.2017.09.028

Ritchie, M. E., Phipson, B., Wu, D., Hu, Y., Law, C. W., Shi, W., & Smyth, G. K. (2015). Limma powers differential expression analyses for RNA-sequencing and microarray studies. Nucleic Acids Research, 43(7), e47. https://doi.org/10.1093/nar/gkv007

Saltz, J., Gupta, R., Hou, L., Kurc, T., Singh, P., Nguyen, V., Samaras, D., Shroyer, K. R., Zhao, T., Batiste, R., Van Arnam, J., Cancer Genome Atlas Research Network, Shmulevich, I., Rao, A. U. K., Lazar, A. J., Sharma, A., & Thorsson, V. (2018). Spatial Organization and Molecular Correlation of Tumor-Infiltrating Lymphocytes Using Deep Learning on Pathology Images. Cell Reports, 23(1), 181–193.e7. https://doi.org/10.1016/j.celrep.2018.03.086

Schoenfelder, S., Furlan-Magaril, M., Mifsud, B., Tavares-Cadete, F., Sugar, R., Javierre, B.-M., Nagano, T., Katsman, Y., Sakthidevi, M., Wingett, S. W., Dimitrova, E., Dimond, A., Edelman, L. B., Elderkin, S., Tabbada, K., Darbo, E., Andrews, S., Herman, B., Higgs, A., … Fraser, P. (2015). The pluripotent regulatory circuitry connecting promoters to their long-range interacting elements. Genome Research, 25(4), 582–597. https://doi.org/10.1101/gr.185272.114

Sharma, P., Hu-Lieskovan, S., Wargo, J. A., & Ribas, A. (2017). Primary, Adaptive, and Acquired Resistance to Cancer Immunotherapy. Cell, 168(4), 707–723. https://doi.org/10.1016/j.cell.2017.01.017

Stunnenberg, H. G., International Human Epigenome Consortium, & Hirst, M. (2016). The International Human Epigenome Consortium: A Blueprint for Scientific Collaboration and Discovery. Cell, 167(5), 1145–1149. https://doi.org/10.1016/j.cell.2016.11.007

Teschendorff, A. E., Breeze, C. E., Zheng, S. C., & Beck, S. (2017). A comparison of reference-based algorithms for correcting cell-type heterogeneity in Epigenome-Wide Association Studies. BMC Bioinformatics, 18(1), 105. https://doi.org/10.1186/s12859-017-1511-5

Tirosh, I., Izar, B., Prakadan, S. M., Wadsworth, M. H., Treacy, D., Trombetta, J. J., Rotem, A., Rodman, C., Lian, C., Murphy, G., Fallahi-Sichani, M., Dutton-Regester, K., Lin, J.-R., Cohen, O., Shah, P., Lu, D., Genshaft, A. S., Hughes, T. K., Ziegler, C. G. K., … Garraway, L. A. (2016). Dissecting the multicellular ecosystem of metastatic melanoma by single-cell RNA-seq. Science (New York, N.Y.), 352(6282), 189–196. https://doi.org/10.1126/science.aad0501

Titus, A. J., Gallimore, R. M., Salas, L. A., & Christensen, B. C. (2017). Cell-type deconvolution from DNA methylation: A review of recent applications. Human Molecular Genetics, 26(R2), R216–R224. https://doi.org/10.1093/hmg/ddx275

Visvader, J. E. (2011). Cells of origin in cancer. Nature, 469(7330), 314–322. https://doi.org/10.1038/nature09781

Wang, Q., Armenia, J., Zhang, C., Penson, A. V., Reznik, E., Zhang, L., Minet, T., Ochoa, A., Gross, B. E., Iacobuzio-Donahue, C. A., Betel, D., Taylor, B. S., Gao, J., & Schultz, N. (2018). Unifying cancer and normal RNA sequencing data from different sources. Scientific Data, 5, 180061. https://doi.org/10.1038/sdata.2018.61

Yoshihara, K., Shahmoradgoli, M., Martínez, E., Vegesna, R., Kim, H., Torres-Garcia, W., Treviño, V., Shen, H., Laird, P. W., Levine, D. A., Carter, S. L., Getz, G., Stemke-Hale, K., Mills, G. B., & Verhaak, R. G. W. (2013). Inferring tumour purity and stromal and immune cell admixture from expression data. Nature Communications, 4, 2612. https://doi.org/10.1038/ncomms3612

Zeng, D., Ye, Z., Wu, J., Zhou, R., Fan, X., Wang, G., Huang, Y., Wu, J., Sun, H., Wang, M., Bin, J., Liao, Y., Li, N., Shi, M., & Liao, W. (2020). Macrophage correlates with immunophenotype and predicts anti-PD-L1 response of urothelial cancer. Theranostics, 10(15), 7002–7014. https://doi.org/10.7150/thno.46176

Ziller, M. J., Hansen, K. D., Meissner, A., & Aryee, M. J. (2015). Coverage recommendations for methylation analysis by whole-genome bisulfite sequencing. Nature Methods, 12(3), 230–232, 1 p following 232. https://doi.org/10.1038/nmeth.3152

Zou, J., Lippert, C., Heckerman, D., Aryee, M., & Listgarten, J. (2014). Epigenome-wide association studies without the need for cell-type composition. Nature Methods, 11(3), 309–311. https://doi.org/10.1038/nmeth.2815

