## Additional File 2 for "GEM-DeCan: Improved tumor immune microenvironment profiling through novel gene expression and DNA methylation signatures predicts immunotherapy response"

#### Supplementary text S1

The deconvolution results presented in this paper were obtained with the GEM-DeCan pipeline, which is shown schematically on Figure S1 representing the workflow for gene expression based deconvolution (**Fig. S1a**) and DNA methylation based deconvolution (**Fig. S1b**).

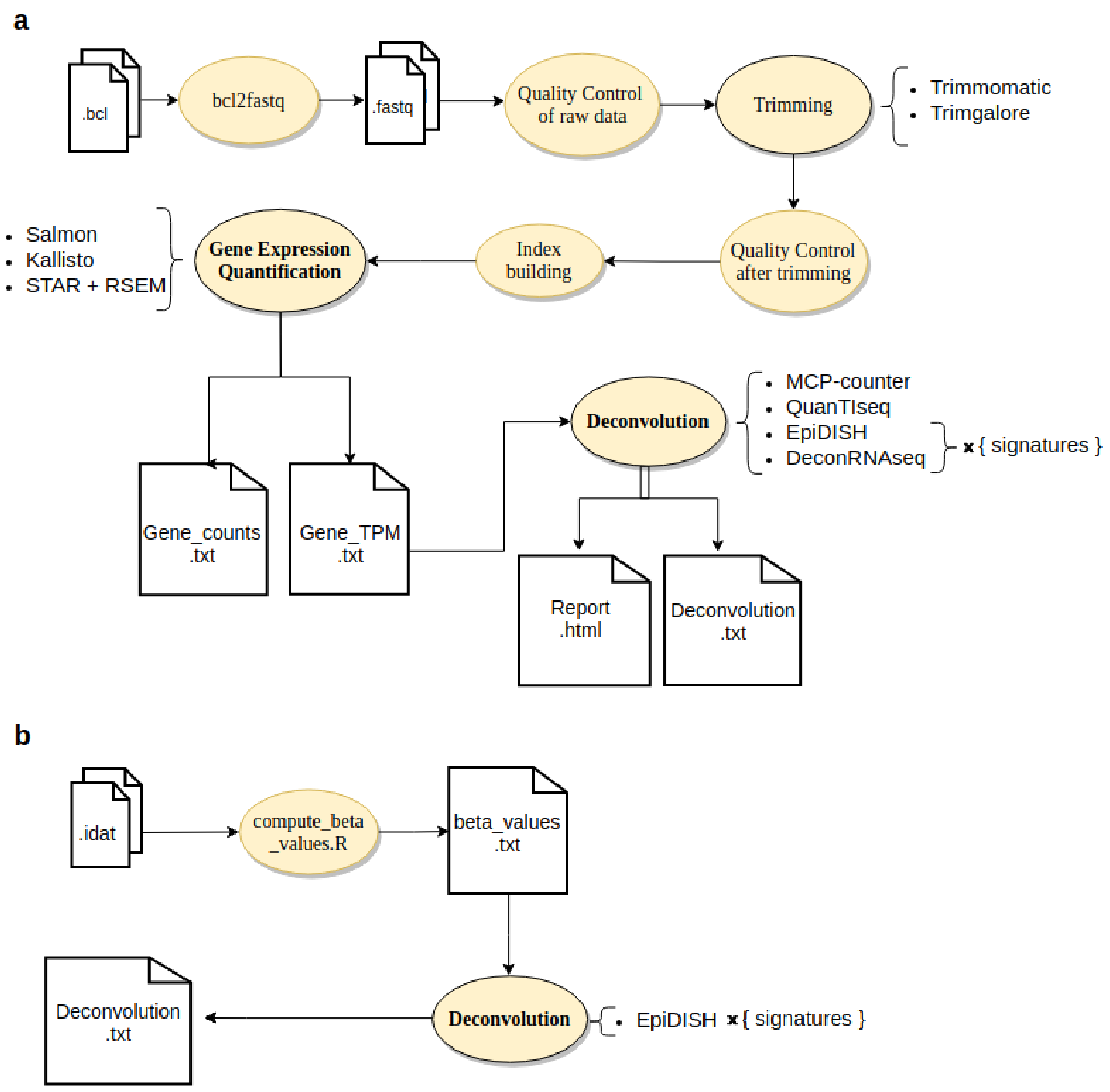


**Fig. S1: Deconvolution pipeline workflow. a)** RNAseq processing and deconvolution pipeline. **b)** DNA methylation deconvolution workflow

#### BPRNA: a signature matrix for immune cell deconvolution in blood

We started by developing a novel immune cell type deconvolution signature matrix based on RNAseq expression data from primary samples for 6 immune cell types including CD4 T cells, CD8 T cells, Monocytes, B cells, NK cells and Neutrophils) in blood, which we called BPRNA (see methods, **Fig. 2b and Additional file 3: Table S1**). We then proceeded to test the BPRNA signature matrix using two deconvolution methods for which signature matrices can be specified by the user (EpiDISH (Teschendorff et al. 2017) and deconRNASeq (Gong 2013)) on peripheral blood mononuclear cell (PBMC) samples ( [Table S1](#)).

#### Testing the BPRNA signature matrix on PBMC samples

We tested our BPRNA signature matrix using PBMC mixtures where both RNA-seq and flow cytometry data was available (Monaco et al. 2019). We observed lower accuracy of estimated total cell fractions for this dataset, especially for NK and CD8 cells (**Fig S2a**). This is probably due to the limited number of samples available for NK (N=2) and to a possible discrepancy between the CD8 activation state in these PBMC samples and the ones used in the signature matrix (which exclude naive CD8 cells).

Multiple gene expression reference-based methods have demonstrated a high accuracy in estimating cell proportions in blood and immune infiltrates from bulk RNA-seq data, amongst which we chose two for comparison to estimates produced by BPRNA: we first compared to two methods that come with their own signature: MCP-counter (Becht et al. 2016), which is a scoring method based on marker genes, and quanTIseq (Finotello & Trajanoski, 2018), which is based on constrained least squares regression and can estimate immune cell fractions and fractions of unknown cells with high accuracy. Finally, we tested the deconRNASeq (Gong & Szustakowski, 2013) method, using different signature matrices, to compare its performance with EpiDISH. The results of running deconvolution with all different methods and signature matrices compared to FACS estimates in PBMC are summarised in **Figure S2b**.

Considering all cell types together or sub-cell types, we observed the combination of BPRNA with EpiDISH compared very favorably relative to the combination of BPRNA with deconRNASeq, which overall showed the weakest performance (**Fig S2b**). However, we note that MCP-counter or quanTIseq outperformed the combination of BPRNA with EpiDISH in several sub-cell types (CD8 T cells, Monocytes, B cells, NK cells, and Neutrophils).

**Figure S2. RNAseq based deconvolution results in PBMC samples. (a)** Pearson Correlation between the cell fractions estimated by EpiDISH using the BPRNA signature vs. FACS proportions in PBMC used as the gold standard (Monaco et al. GSE107011). (**b**) Pearson correlation of estimates using different methods and FACS gold standard. The significance of the Pearson Correlation is indicated by stars: *p<0.05, **p<0.01, ***p<0.001. Note that the MCP_counter signature doesn't include CD4.

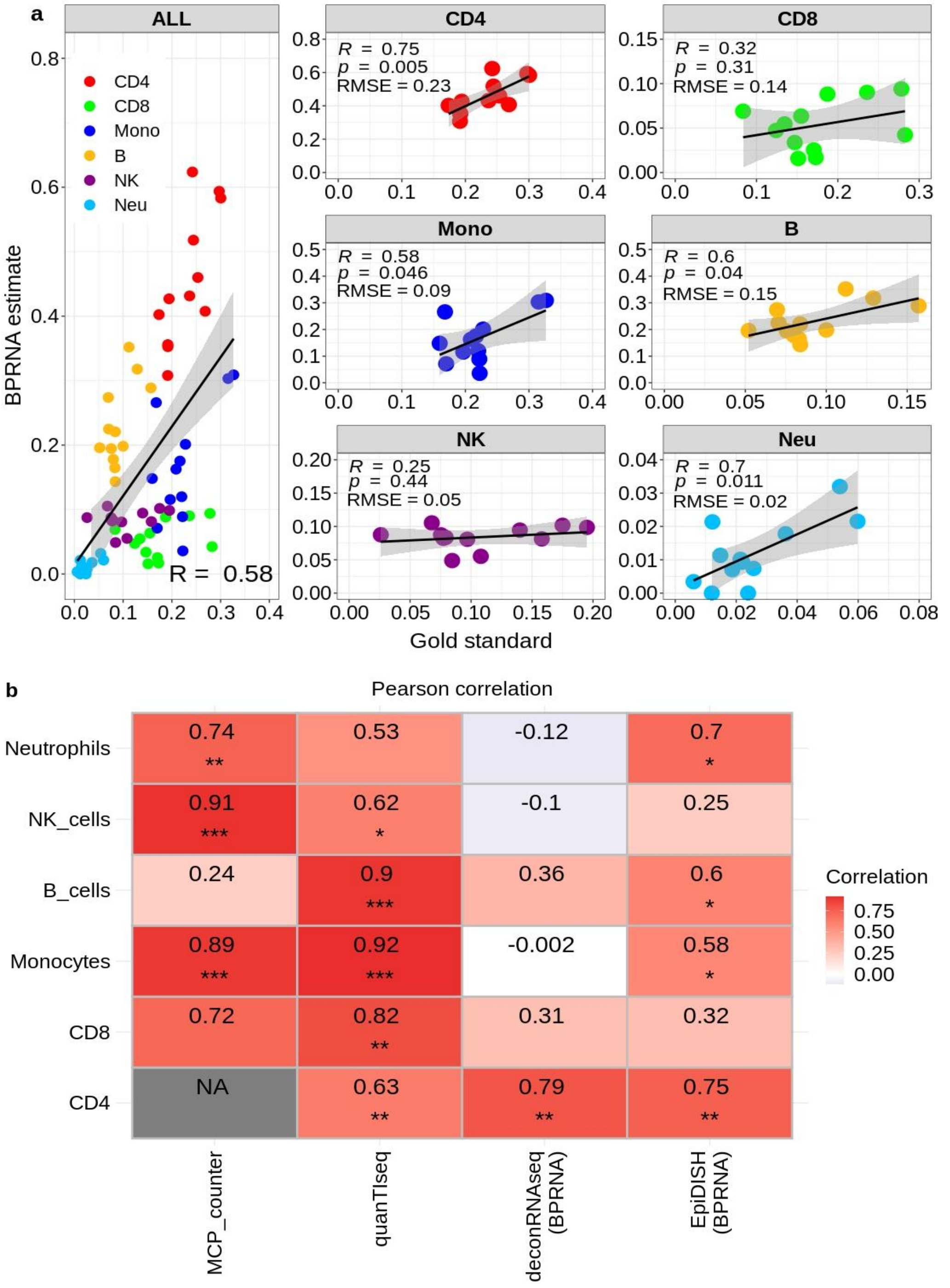

### Supplementary text S2

#### Testing the CCLE_TIL10 and BPRNACan signatures on in-silico immune and cancer cell line mixtures

We started with a set of in-silico samples in which cell type proportions are known. This RNAseq dataset (see **Table S1**) is composed of reads from purified samples from 10 immune cells (B-cells, NK-cells, CD4+ T-cells, CD8+ T-cells, Monocytes, NK, Neutrophils, M1, M2 and Dendritic cells), which were added to reads from a sample of MCF10 cancer cell lines in different proportions (Finotello & Trajanoski, 2018). The results of using the CCLE_TIL10 signature matrix on this in-silico mixture are in excellent agreement with true mRNA proportions (Pearson R > 0.9), for 11 cell types (**Figure S3a**), as expected, since the TIL10 signature matrix was derived from this data and the CCLE_TIL10 signature matrix is an extension of it.

The proportions estimated using our newly developed BPRNACan signature matrix (with the EpiDISH method) were in very good agreement with the true proportions (Pearson R = 0.96 for cancer cells and Pearson R > 0.74 for other immune types, **Figure S3b**), except for macrophages M2 (R = 0.35). This lower performance in the detection of M2 macrophages could be due to BPRNACan missing M2s in some samples, suggesting that the signature matrix potentially does not capture all M2 phenotypes that are present. Our signature matrix might also have similar issues with M1 macrophages, as we can clearly see two groups of samples, one of which has estimated M1 proportions quite discordant from the true cell fractions (**Figure S3b**).

**Figure S3.** Correlation between estimated fractions from (**a**) CCLE_TIL10 and (**b**) BPRNACan signatures using EpiDISH and true mRNA fraction (Finotello et al. 2019) for each cell subtype in known -proportion in-silico mixture (Dataset 6).

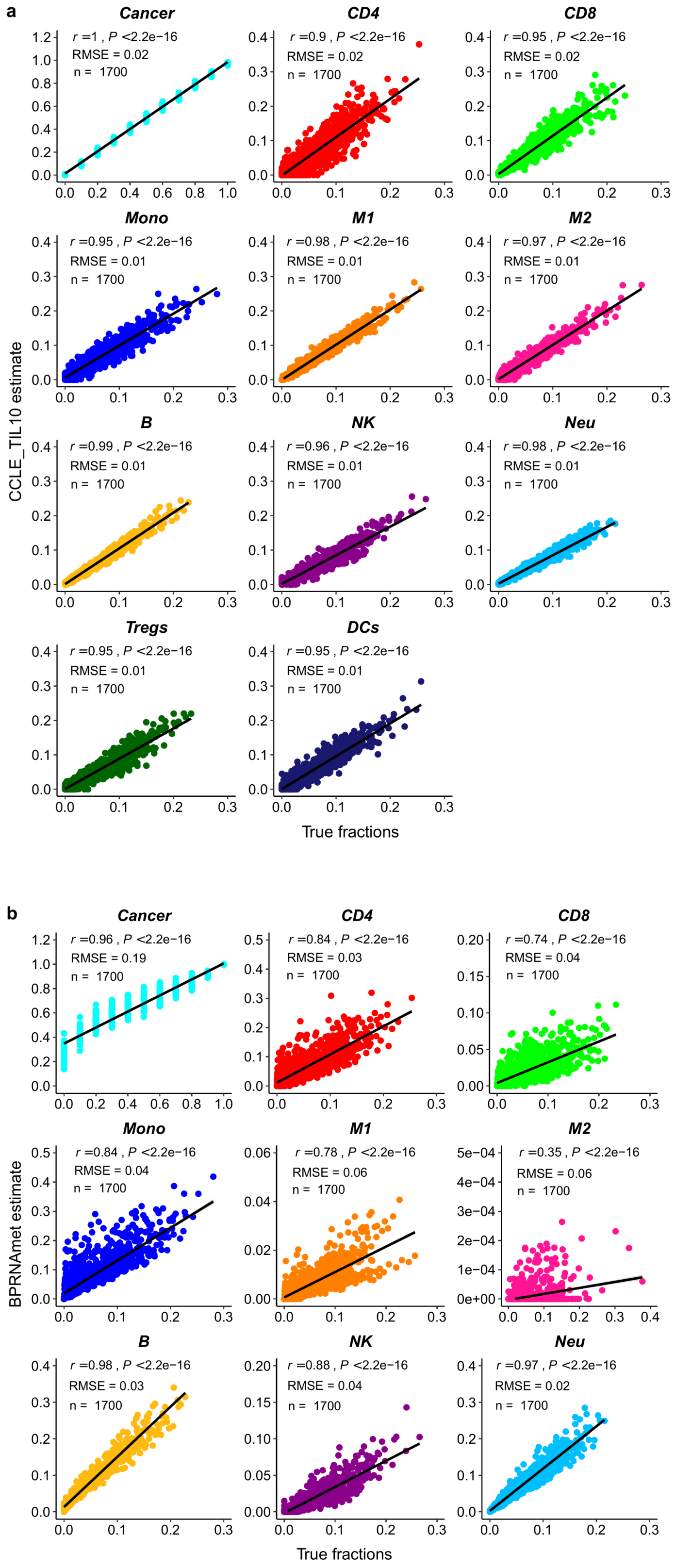

We then compared estimates of tumor purity using CCLE_TIL10 (**Figure S4a**) and BPRNACan estimates (**Figure S4b**).

**Figure S4.** Comparing TCGA-LUAD tumor purity proportion using CCLE_TIL10 and BPRNACan with other previously published purity estimation methods by Pearson correlations. (**a**) Predicted tumor purity between CCLE_TIL10 and previously purity estimation methods. (**b**) The scatter plots of tumor purities predicted by BPRNACan and the available purity estimation methods.

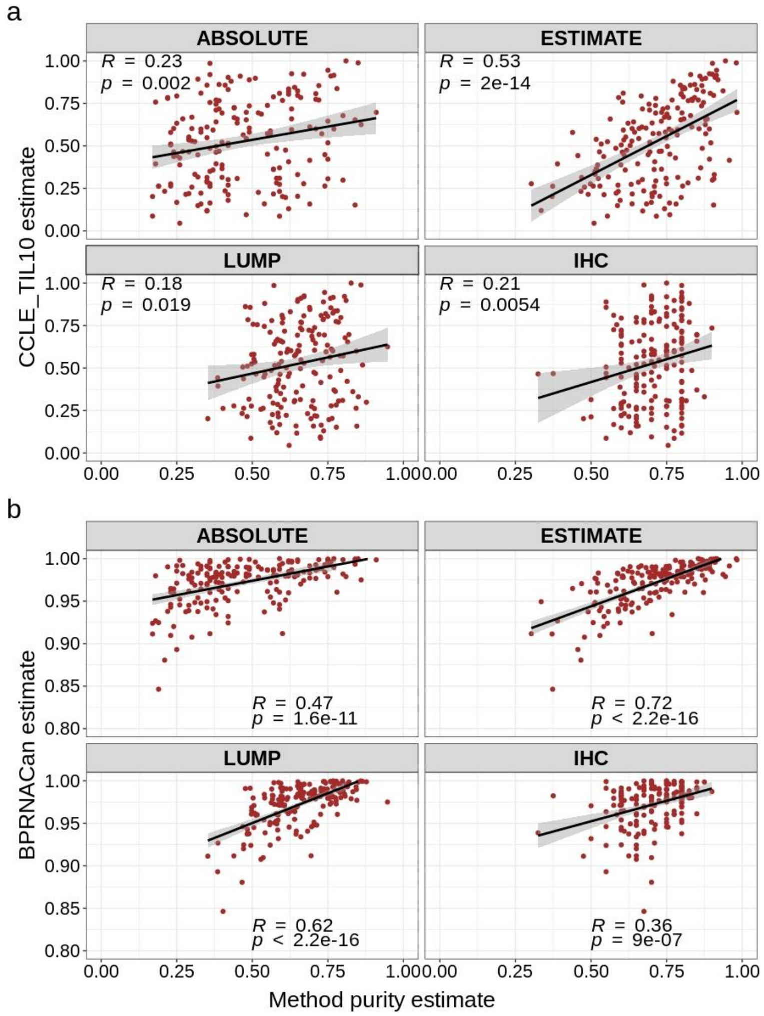

### Supplementary text S3

#### Testing the BPmet signature matrix on peripheral blood samples

To evaluate the performance of the BPmet signature matrix using the EpiDISH method, we first applied it to an independent publicly available Illumina EPIC array (850k CpGs) dataset choosing 100 whole blood samples from healthy donors with the true cell composition verified by FACS (gold standard) (GSE132203, Grady Trauma Project). We observed extremely high Pearson correlations between the estimated cell compositions and FACS fractions for all included samples (Pearson's R = 0.993, $p < 2.2e{-16}$) and each cell subtype (**Figure S5a**).

We then compared the results from using the BPmet signature matrix in these 100 whole blood samples from the previously mentioned gold standard dataset (GSE132203) to the results using other signature matrices. The signature matrices we compared with the MethylCIBERSORT signature matrix, and the default signature matrix for the EpiDISH method (Zheng et al., 2018), based on DNAse hypersensitivity sites (DHS), which are highly cell-type specific regions of open chromatin. Altogether, correlations between the different signature matrix estimates and the FACS fractions were similar for EpiDISH-DHS and BPmet, while MethylCIBERSORT had a worse performance. Only NK cells estimate was worse correlated with BPmet compared to EpiDISH-DHS and MethylCIBERSORT, probably due to the few samples included for this cell type in creating our signature matrix (NK: n = 2) (**Figure S5b,c**)

**Figure S5.** Comparing different DNA methylation signatures on 100 whole blood data with FACS (GSE132203) from the Grady Trauma Project using the EpiDISH RPC method. (**a**) BPmet, (**b**) MethylCIBERSORT, (**c**) EPIDISH-DHS

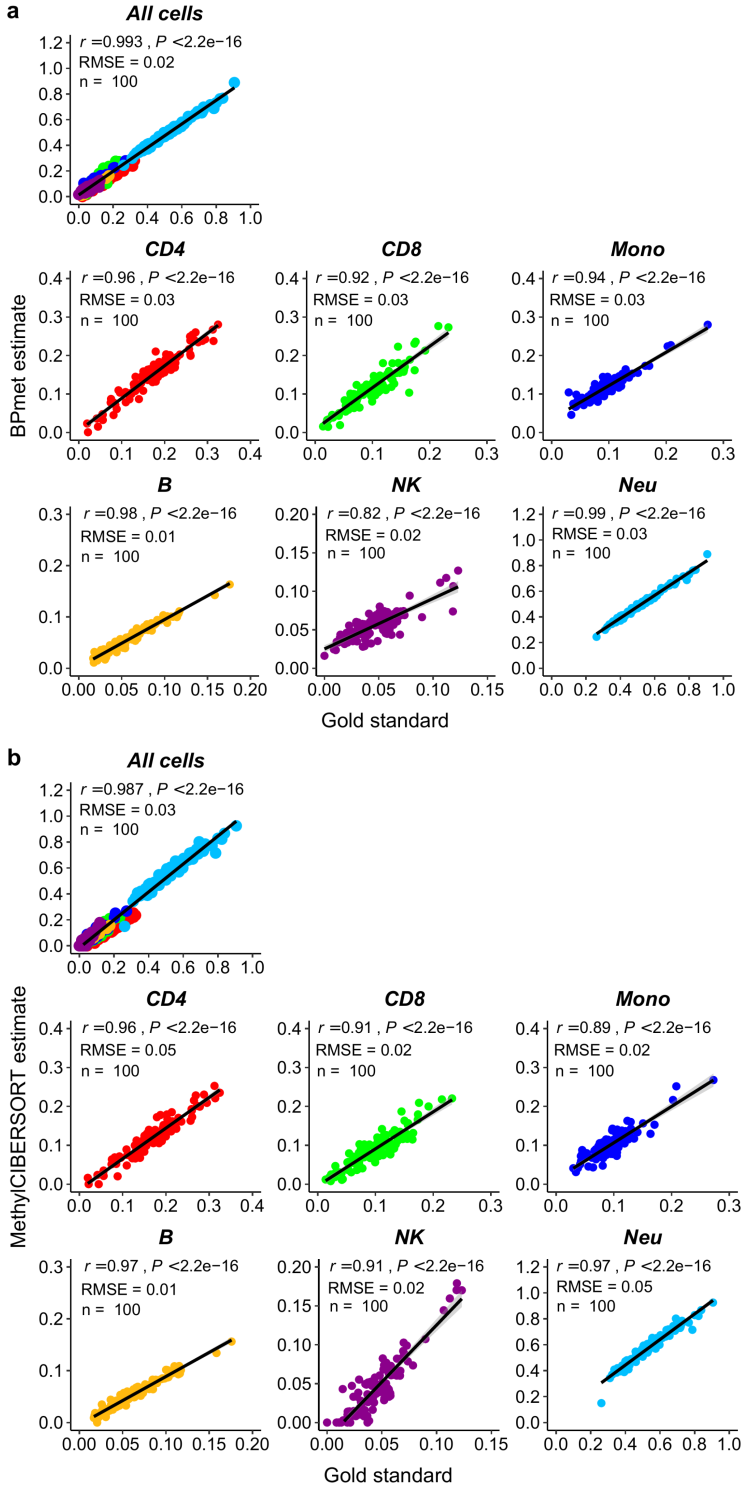

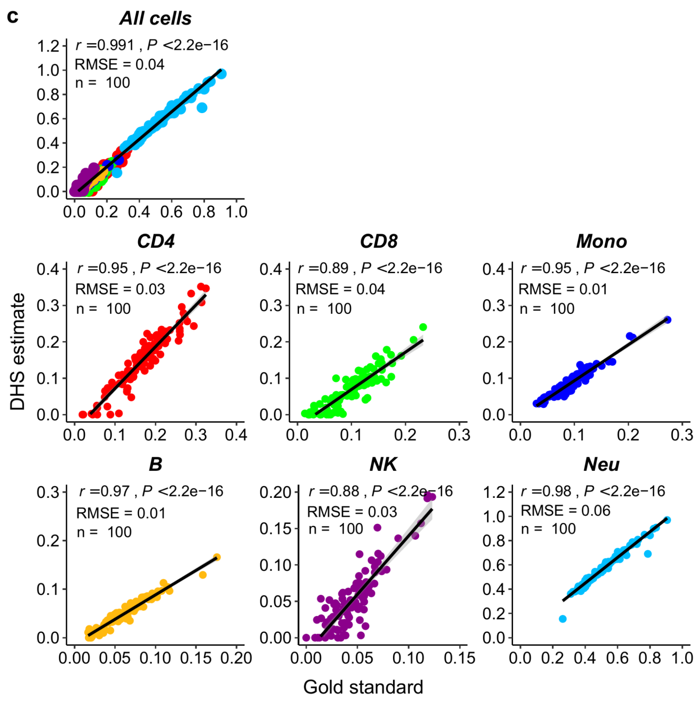


Since the MethylCIBERSORT and EpiDISH-DHS signature matrices were generated starting from the Illumina 450k platform, we tested them on another set of 6 whole blood samples analysed by the same technology and compared results to the flow cytometry measurements provided for that dataset [32]. We were able to obtain better correlations (Pearson's R = 0.93, P < 2.2e-16) for all cell types confounded compared to either methylCIBERSORT or EpiDISH-DHS. We obtained correlations above 0.94 for each cell subtype except for NK cells (**Fig. 3b and Additional file 2: Figure S6**). We thus conclude that our BPmet signature matrix can correctly capture cell type composition as well or better than other available methods in whole blood.

**Figure S6.** Comparing different DNA methylation signatures in 6 WB data analysed by illumina 450k platform with FACS (GSE77797) [1] (**a**) BPmet (**b**) MethylCIBERSORT (**c**) DHS (hypersensitivity sites)

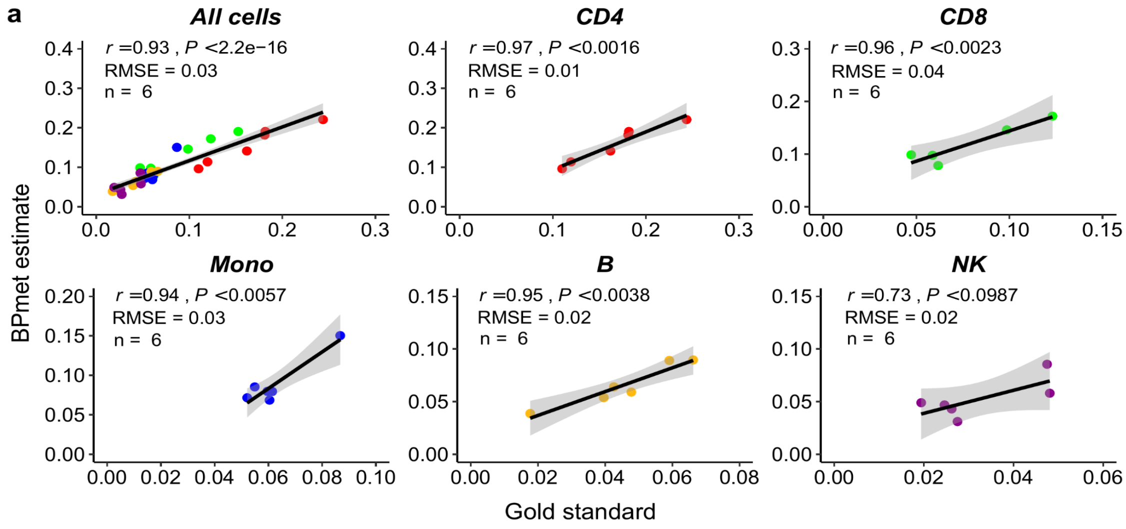

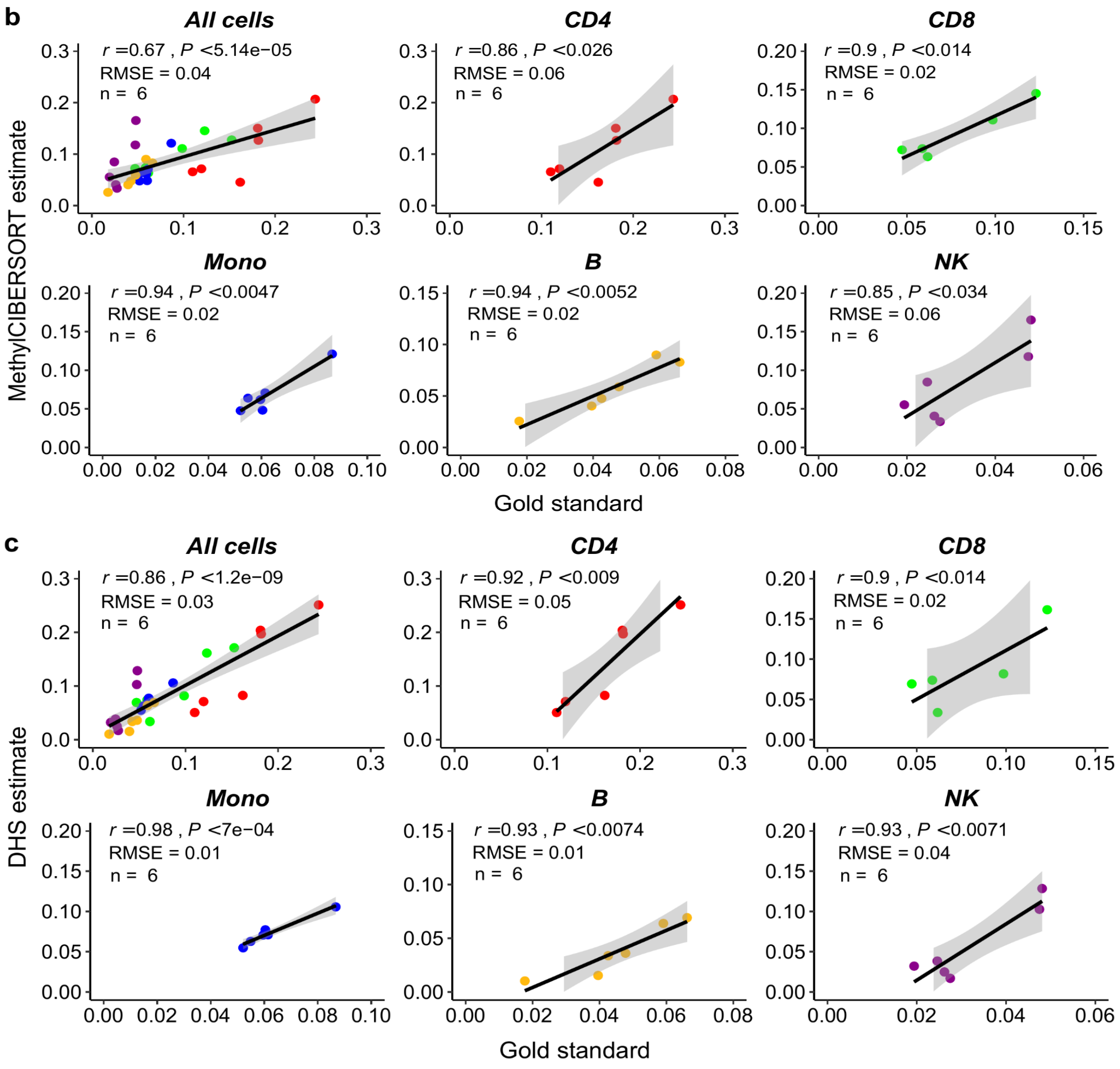

b
All cells
r =0.67 , P <5.14e−05
RMSE = 0.04
n = 6
CD4
r =0.86 , P <0.026
RMSE = 0.06
n = 6
CD8
r =0.9 , P <0.014
RMSE = 0.02
n = 6
Mono
r =0.94 , P <0.0047
RMSE = 0.02
n = 6
B
r =0.94 , P <0.0052
RMSE = 0.02
n = 6
NK
r =0.85 , P <0.034
RMSE = 0.06
n = 6
MethylCIBERSORT estimate
Gold standard
c
All cells
r =0.86 , P <1.2e−09
RMSE = 0.03
n = 6
CD4
r =0.92 , P <0.009
RMSE = 0.05
n = 6
CD8
r =0.9 , P <0.014
RMSE = 0.02
n = 6
Mono
r =0.98 , P <7e−04
RMSE = 0.01
n = 6
B
r =0.93 , P <0.0074
RMSE = 0.01
n = 6
NK
r =0.93 , P <0.0071
RMSE = 0.04
n = 6
DHS estimate
Gold standard

We then turned to measuring performance of our DNA methylation based deconvolution on tumour samples from TCGA

**Figure S7: Deconvolution using different DNA methylation signature matrices in tumor samples.** (**a**) Correlations between TCGA-LUAD tumor purity proportion estimated by BPmetCan and other previously published purity estimation methods compared to estimates by ABSOLUTE using Pearson correlations [12]. (**b**) Scatterplots showing correlation between estimates of proportions of different immune cell types and H&E image estimates from [50].

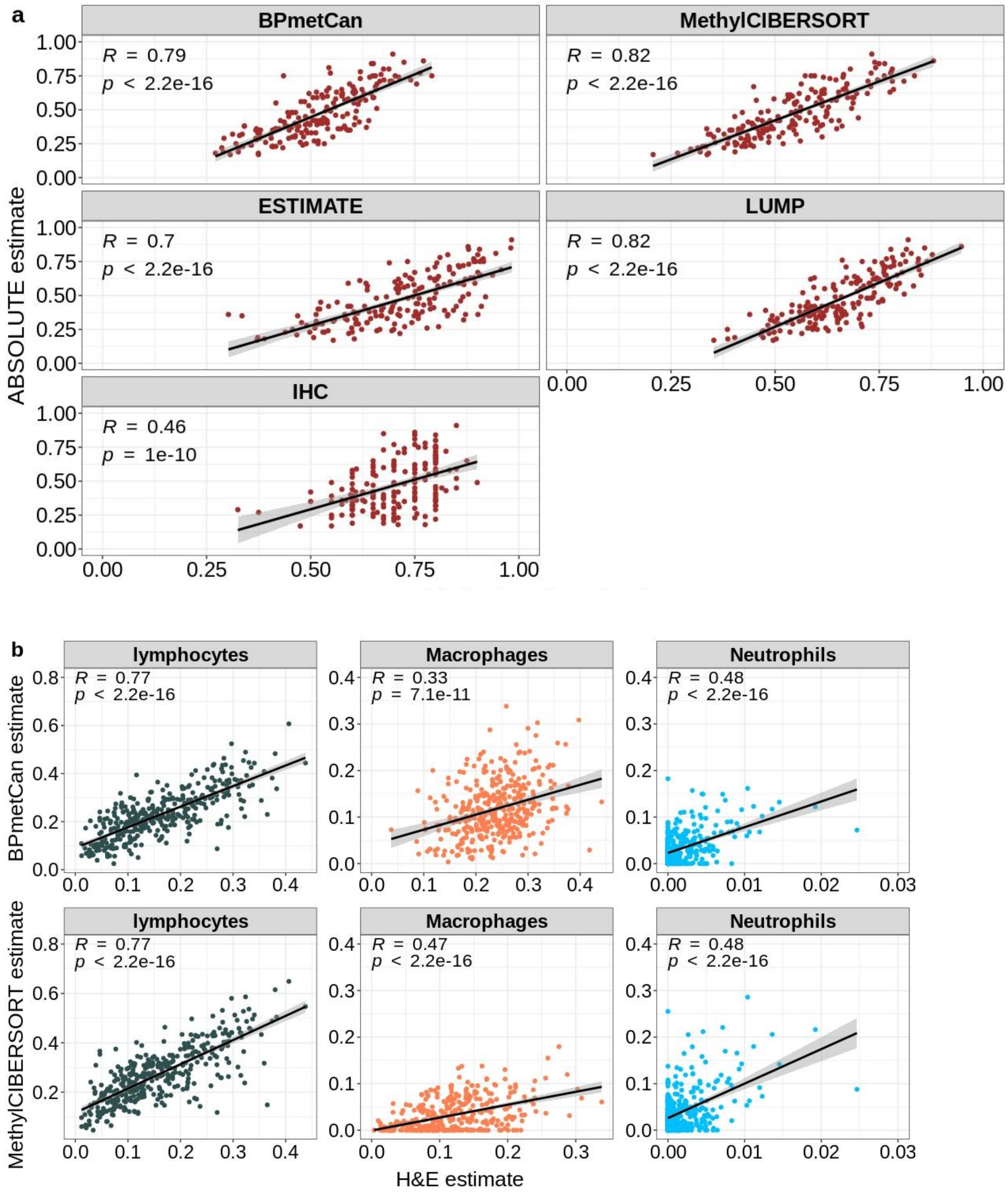

#### Supplementary Tables

**Table S1.** Main datasets used to evaluate our signature in this study: data set ID, sample type and size, availability of deconvolution, selection of validation data, Platform, reference publication.

| ID | Samples | Deconvolution | Selected samples | Platform | Reference |
|---|---|---|---|---|---|
| 1 | 795 DNA methylation was assessed in whole blood from participants of the Grady Trauma Project | DNA methylation | First 100 whole blood samples with Flow Cytometry estimates: $CD4^+$ and $CD8^+$ T cells, monocytes, B cells, NK cells, neutrophils (GSE132203) | Illumina EPIC | GSE132203 |
| 2 | 12 Reconstructed whole blood mixtures with other 6 whole blood samples with Flow Cytometry estimates | DNA methylation | 6 whole blood samples with Flow Cytometry estimates: $CD4^+$ and $CD8^+$ T cells, monocytes, B cells, NK cells. (GSE77797) | Illumina 450k | [4] |
| 3 | 21 TCGA cancer types | DNA methylation / RNAseq | 180 Lung Adenocarcinoma without missing values / 489 Breast Invasive Carcinoma no missing values with other 3 cancer purity estimated methods | Illumina 450k / RNAseq | [5] |
| 4 | 13 TCGA cancer types | DNA methylation / RNAseq | 413 Lung Adenocarcinoma lymphocytes, macrophages and neutrophils scores computed from H&E stained section slides | Images | [6] |
| 5 | Total RNA of 29 immune cell types (from 4 individuals) and 13 individuals peripheral blood mononuclear cells (PBMCs) | RNAseq | 13 individuals PBMCs with Flow Cytometry estimates: $CD4^+$ and $CD8^+$ T cells, monocytes, B cells, NK cells (GSE107011) | illumina RNAseq | [2] |
| 6 | 1700 simulated RNA-seq data sets with various immune cell fractions and tumor contents (http://icbi.at/quantiseq). | RNAseq | 1700 In-silico RNAseq with true mRNA compositions: $CD4^+$ and $CD8^+$ T cells, monocytes, M1 and M2 macrophages, B cells, NK cells, neutrophils, $T_{reg}$ cells, and dendritic cells (http://icbi.at/quantiseq) | Illumina RNAseq | [3] |
| 7 | 59 multiple myeloma samples, generated by removal of cancer cells from bone marrow samples | RNAseq | Non-tumoral of 59 multiple myeloma samples with Flow Cytometry estimates of $CD4^+$ and $CD8^+$ T cells and NK cells | Illumina RNAseq | [7] |
| 8 | Single-cell RNAseq from 19 donors and comprises primary tumors, lymph node metastasis or other lesions. | Single-cell RNAseq | 19 reconstructed non metastasis bulk samples from each donor with true mRNA cell fractions of Cancer, $CD4^+$ and $CD8^+$ T cells, Macrophage, B cells, NK cells and Tregs (GSE72056) | Illumina NextSeq | [9] |
| 9 | RNA-seq from lymph node bulk samples from 4 melanoma patients | RNAseq | 4 melanoma patients form lymph node bulk RNA-seq with Flow Cytometry estimates of Cancer, $CD4^+$ and $CD8^+$ T cells, B cells and NK cells (GSE93722) | Illumina RNAseq | [8] |

**Table S2:** Main reference-based deconvolution methods used: Methods, availability of deconvolution, availability of reference profiles, reference publication.

| ID | Methods | Deconvolution | Cell estimates | Reference |
|---|---|---|---|---|
| 1 | EpiDISH | DNAm / RNAseq | Original method: $CD4^+$ and $CD8^+$ T cells, monocytes, B cells, NK cells, neutrophils, Eosinophils. This method can be used with alternative signatures increasing number of cell types quantified | [10] |
| 2 | MethylCIBERSORT | DNAm | Cancer, $CD4^+$ and $CD8^+$ T cells, monocytes, B cells, NK cells, neutrophils, $T_{reg}$ cells, Eosinophils, Endothelial, Fibroblast | [11] |
| 3 | DeconRNAseq | RNAseq | Any | [12] |
| 4 | MCP-counter | RNAseq | T cells, $CD8^+$ T cells, Monocytic lineage, B lineage,NK cells, Neutrophils, Endothelial cells, Fibroblasts, Cytotoxic lymphocytes, Myeloid dendritic cells, | [13] |
| 5 | quanTIseq | RNAseq | $CD4^+$ and $CD8^+$ T cells, monocytes, M1 and M2 macrophages, B cells, NK cells, neutrophils, $T_{reg}$ cells, and dendritic cells | [3] |
| 6 | CIBERSORTx | RNAseq | Can be used with any signature derived from scRNAseq data. We used 3 different signatures provided in the original software (Fig2ab-NSCLC_PBMCs_scRNAseq_sigmatrix, sigmatrix_HNSCC_Fig2cd, mixture_HNSCC_Puram_et_al_Fig2cd ) and also our own. | [https://www.nature.com/articles/s41587-019-0114-2] |
| 7 | ABSOLUTE | DNAm / RNAseq | Involves analysing somatic DNA alterations to estimate ploidy and hence tumour purity | [14] |
| 8 | ESTIMATE | DNAm / RNAseq | Based on single sample Gene Set Enrichment Analysis for tumour purity | [15] |
| 9 | LUMP & IHC | DNAm / RNAseq | Based on averaging the 44 unmethylated immune-specific CpG sites or immunohistochemistry for tumour purity | [5] |

**Table S3.** Fisher's tests

**a**

| **Illumina annotation** | In Gene signature | Not in Gene Signature |
|---|---|---|
| Genes with cpgs in promoter | 24 | 145 |
| Genes without cpgs in promoter | 1364 | 24506 |

p.value = 1.138e-05

**b**

| **Blueprint annotation** | In Gene signature | Not in Gene Signature |
|---|---|---|
| Genes with cpgs in promoter | 52 | 594 |
| Genes without cpgs in promoter | 1322 | 36538 |

p.value = 7.34e-08

**Table S4.** PCHi-C network analysis for BPmetCan signature

| | | |
|---|---|---|
| | Total nodes in network (BP all) | 249511 |
| | Promoter nodes in network | 20582 |
| | Oe nodes in network | 228 929 |
| | Promoter nodes in BPRNACan gene sign | 1127 |
| | Total nodes in CpG sign | 1131 |
| BPMetPro | Nodes in CpG sign also prom | 348 (645 unique genes) |
| BPMetOe | Nodes in CpG sign also oe | 779 |
| BPRNAMetSig | Nodes in CpG sign prom and in gene sign | 64 (52 uniques genes) |
| BP3DMet | prom neighbours of CpG sign nodes | 3642 (5478 uniques genes) |
| | prom neighbours of CpG sign nodes also in gene sig | 255 |
| | prom neighbours of the 64 CpGs sign prom and in gene | 630 (76 from gene sig, 554 other genes) |

| | | |
|---|---|---|
| | sign | |
| | prom neighbours of non prom CpGs sign | 2716 (132 from gene sig, 2584 other genes) |
| BPRNACan | Total genes in gene sign | 1403 |
| BPmetCan | Total CpGs in CpG sig | 1896 |
| BPMetPro - BPRNAMetSig (BPRNAProMet) | Genes in BPmetCan with CpGs associated to a promoter, but not present in BPRNACan | 593 |

#### References


1. Koestler DC, Jones MJ, Usset J, Christensen BC, Butler RA, Kobor MS, et al. Improving cell mixture deconvolution by identifying optimal DNA methylation libraries (IDOL) [Internet]. BMC Bioinformatics. 2016. Available from: http://dx.doi.org/10.1186/s12859-016-0943-7

2. Monaco G, Lee B, Xu W, Mustafah S, Hwang YY, Carré C, et al. RNA-Seq Signatures Normalized by mRNA Abundance Allow Absolute Deconvolution of Human Immune Cell Types. Cell Rep. 2019;26:1627–40.e7.

3. Finotello F, Mayer C, Plattner C, Laschober G, Rieder D, Hackl H, et al. Molecular and pharmacological modulators of the tumor immune contexture revealed by deconvolution of RNA-seq data. Genome Med. 2019;11:34.

4. Koestler DC, Jones MJ, Usset J, Christensen BC, Butler RA, Kobor MS, et al. Improving cell mixture deconvolution by identifying optimal DNA methylation libraries (IDOL) [Internet]. BMC Bioinformatics. 2016. Available from: http://dx.doi.org/10.1186/s12859-016-0943-7

5. Aran D, Sirota M, Butte AJ. Systematic pan-cancer analysis of tumour purity. Nat Commun. 2015;6:8971.

6. Saltz J, Gupta R, Hou L, Kurc T, Singh P, Nguyen V, et al. Spatial Organization and Molecular Correlation of Tumor-Infiltrating Lymphocytes Using Deep Learning on Pathology Images. Cell Rep. 2018;23:181–93.e7.

7. Nakamura K, Kassem S, Cleynen A, Chrétien M-L, Guillerey C, Putz EM, et al. Dysregulated IL-18 Is a Key Driver of Immunosuppression and a Possible Therapeutic Target in the Multiple Myeloma Microenvironment [Internet]. Cancer Cell. 2018. p. 634–48.e5. Available from: http://dx.doi.org/10.1016/j.ccell.2018.02.007

8. Racle J, de Jonge K, Baumgaertner P, Speiser DE, Gfeller D. Simultaneous enumeration of cancer and immune cell types from bulk tumor gene expression data. Elife [Internet]. 2017;6. Available from: http://dx.doi.org/10.7554/eLife.26476

9. Tirosh I, Izar B, Prakadan SM, Wadsworth MH 2nd, Treacy D, Trombetta JJ, et al. Dissecting the multicellular ecosystem of metastatic melanoma by single-cell RNA-seq. Science. 2016;352:189–96.

10. Teschendorff AE, Breeze CE, Zheng SC, Beck S. A comparison of reference-based algorithms for correcting cell-type heterogeneity in Epigenome-Wide Association Studies. BMC Bioinformatics. 2017;18:105.

11. Chakravarthy A, Furness A, Joshi K, Ghorani E, Ford K, Ward MJ, et al. Author Correction:

Pan-cancer deconvolution of tumour composition using DNA methylation. Nat Commun. 2018;9:4642.

12. Gong T, Szustakowski JD. DeconRNASeq: a statistical framework for deconvolution of heterogeneous tissue samples based on mRNA-Seq data. Bioinformatics. 2013;29:1083–5.

13. Becht E, Giraldo NA, Lacroix L, Buttard B, Elarouci N, Petitprez F, et al. Estimating the population abundance of tissue-infiltrating immune and stromal cell populations using gene expression. Genome Biol. 2016;17:218.

14. Carter SL, Cibulskis K, Helman E, McKenna A, Shen H, Zack T, et al. Absolute quantification of somatic DNA alterations in human cancer [Internet]. Nature Biotechnology. 2012. p. 413–21. Available from: http://dx.doi.org/10.1038/nbt.2203

15. Yoshihara K, Shahmoradgoli M, Martínez E, Vegesna R, Kim H, Torres-Garcia W, et al. Inferring tumour purity and stromal and immune cell admixture from expression data. Nat Commun. 2013;4:2612.
